# Single cell transcriptomics identifies a unique adipocyte population that regulates bone marrow environment

**DOI:** 10.1101/754481

**Authors:** Leilei Zhong, Lutian Yao, Robert J. Tower, Yulong Wei, Zhen Miao, Jihwan Park, Rojesh Shrestha, Luqiang Wang, Wei Yu, Nicholas Holdreith, Yejia Zhang, Wei Tong, Yanqing Gong, Fanxin Long, Jaimo Ahn, Patrick Seale, Katalin Susztak, Mingyao Li, Chider Chen, Ling Qin

## Abstract

Bone marrow mesenchymal lineage cells are a heterogeneous cell population involved in bone homeostasis and diseases such as osteoporosis. While it is long postulated that they originate from mesenchymal stem cells (MSCs), the true identity of MSCs and their in vivo bifurcated differentiation routes into osteoblasts and adipocytes remain poorly understood. Here, by employing single cell transcriptome analysis, we identified MSCs and delineated their bi-lineage differentiation paths in young, adult and aging mice. Among several newly discovered mesenchymal subpopulations, one is a distinct population of adipose-lineage cells that we named marrow environment regulating adipose cells (MERAs). MERAs are non-proliferative, post-progenitor cells that express many mature adipocyte markers but are devoid of lipid droplets. They are abundant in the bone marrow of young mice, acting as pericytes and stromal cells that form numerous connections among themselves and with other cells inside bone, including endothelial cells. Genetic ablation of MERAs disrupts marrow vessel structure, promotes de novo bone formation. Taken together, MERAs represent a unique population of adipose lineage cells that exist only in the bone marrow with critical roles in regulating bone and vessel homeostasis.

---

Osteoporosis is a silently progressive disease characterized by excessive bone loss and structural deterioration until it clinically presents as bone fragility and fracture. While mostly afflicting post-menopausal women and the elderly ^1^, it also manifests in adolescents and adults as a result of endocrine and metabolic abnormalities or medications ^2^. Osteoporosis is largely or partially caused by diminished bone forming activity and often accompanied by increases in marrow adiposity. Therefore, understanding the nature of bone marrow mesenchymal stem cells (MSCs) and their differentiation routes into osteoblasts and adipocytes has always been the center of bone research. However, in contrast to the wealth of knowledge regarding hematopoiesis from hematopoietic stem cells (HSCs) ^3^, there remains a paucity of in vivo knowledge of MSCs and their descendants, which has greatly limited advances in treating clinical disorders of bone loss.

It is well-accepted that bone marrow mesenchymal progenitors are heterogeneous, including MSCs and their descendants at various differentiation stages before they reach the terminal states as osteoblasts, osteocytes, and adipocytes. Our current knowledge of those progenitors is mostly dependent on the in vitro expansion for CFU-F assay, multi-lineage differentiation assays, and in vivo transplantation. However, in vitro culture does not necessarily replicate in vivo cell behavior after depleting their environmental cues. Another common approach is lineage tracing that uses a specific promoter-driven *Cre* or *CreER* system to label a portion of mesenchymal lineage cells. However, it generally does not provide information about the specific stage(s) of mesenchymal progenitors that cells start to be labeled. The recently available large-scale single cell RNA-sequencing (scRNA-seq), which is capable of identifying and interrogating rare cell populations and deducting the course of differentiation ^4^, finally provides us an unbiased tool to investigate bone marrow mesenchymal cells in vivo.

Three reports published in the past few months applied this technique on mouse bone marrow mesenchymal cells. Based on previous studies that LepR marks adult bone marrow MSCs ^5^ and LepR^+^ cells serve as niche for hematopoietic progenitors ^6^, one study used *Lepr-Cre* to label mesenchymal stromal cells and *Col2.3-Cre* to label osteoblasts in order to analyze HSC niches ^7^. The other two studies depleted hematopoietic cells from bone marrow to analyze the remaining bone marrow cells ^8, 9^. Interestingly, all those studies identified a large MSC cluster expressing many adipocyte markers. For our study, we used a different approach by taking advantage of a *Col2-Cre Rosa-tdTomato (Col2/Td)* mouse model that we and others previously reported to label bone marrow mesenchymal lineage cells ^10, 11^. Since in this mouse model, Td labels every osteocyte and every bone marrow adipocyte in vivo and almost all CFU-F colonies derived from bone marrow, we reason that all bone marrow mesenchymal lineage cells, including MSCs and their progeny progenitors, are included within the Td^+^ population. By applying large scale scRNA-seq on this population derived from various age groups, we identified an in vivo MSC population different from the other three reports ^7–9^ and delineated the in vivo evolvement of MSCs into mature osteogenic and adipogenic cells through hierarchical differentiation paths. Among newly identified mesenchymal subpopulations, a new type of adipose lineage cells is subsequently validated and investigated for their critical actions in regulating bone marrow vasculature and bone homeostasis under normal and injury conditions.

## Results

### Single cell transcriptomic profiling of bone marrow mesenchymal lineage cells

In 1-month-old *Col2/Td* mice, in line with our previous report ^11^, Td labeled all growth plate chondrocytes (1052 out of 1052 chondrocytes counted, n=4 mice, 100%), osteoblasts, osteocytes (3600 out of 3600 osteocytes counted, n=4 mice, 100%), CD45^−^ stromal cells (2050 out of 2050 cells with stromal morphology counted, n=4 mice, 100%), Perilipin^+^ adipocytes (5 out of 5 adipocytes counted, n=4 mice, 100%), and pericytes (Fig. 1A, S1A). Td^+^ stromal cells displayed reticular shapes and did not overlap with hematopoietic (CD45^+^) or endothelial (Emcn^+^) cells. We previously developed an enzymatic digestion method to release bone marrow cells trapped within the trabecular bone (endosteal bone marrow) and demonstrated that they contain a higher frequency of mesenchymal progenitors than central bone marrow flushed from the diaphysis of long bones ^12^. This enzymatic digestion method collects osteoblasts as well and thus, is better than conventional flushing method by providing a definite cluster of mature cells for single cell analysis. Due to its small quantity, Td^+^ peak among endosteal bone marrow cells was sometimes distinguishable (1.2±0.1%) but often not. Nevertheless, fractioning all cells into the top 1%, 1-2%, and >2% cells based on the Td signal (Fig. S1B) revealed that the top 1% group contains all the CFU-F forming cells (Fig. S1C) and almost all CFU-F colonies (98.7%) are Td^+^ (Fig. S1D). Since osteoblasts are also Td^+^, these data suggested that the top 1% cells include all bone marrow mesenchymal progenitors as well as osteoblasts.

**Figure 1.**
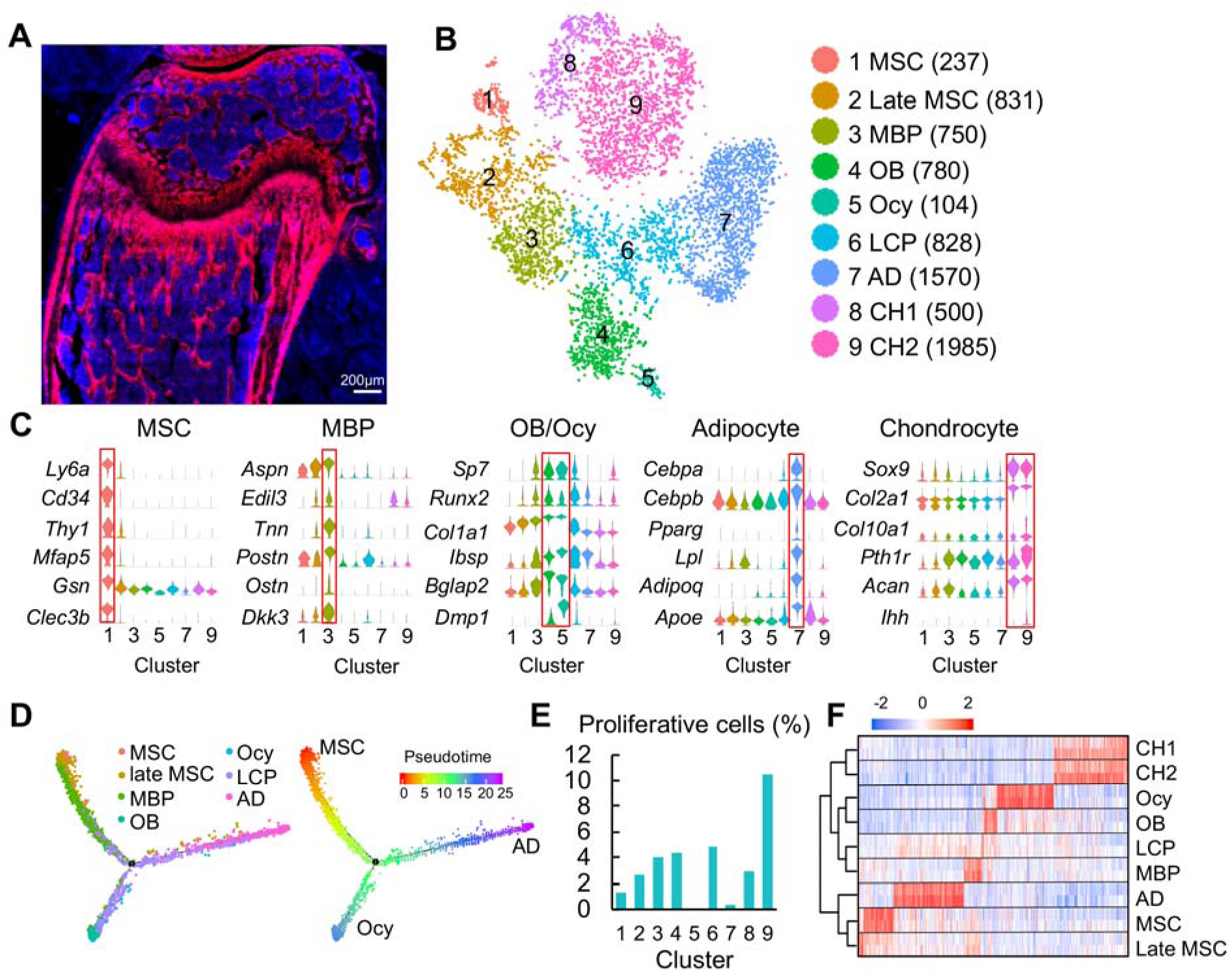
Clustering of bone marrow mesenchymal lineage cells by single cell transcriptomics reveals the in vivo identity of novel mesenchymal subpopulations. (A) Fluorescent image of distal femur of 1-month-old *Col2/Td* mice. (B) The tSNE plot of 7585 Td^+^ mesenchymal lineage cells isolated from endosteal bone marrow of 1-1.5-month-old *Col2/Td* mice (n=5). Cell numbers are listed in parenthesis next to cluster names. MBP: mesenchymal bi-potent progenitor; OB: osteoblast; Ocy: osteocyte; LCP: lineage committed progenitor; AD: adipocyte; CH: chondrocyte. (C) Violin plots of marker gene expression for MSC, MBP, osteoblast/osteocyte, adipocyte, and chondrocyte clusters. (D) Monocle trajectory plot of bone marrow mesenchymal lineage cells. Cells are labeled according to their Seurat clusters. Pseudotime scale is shown on the right. (E) The percentage of proliferative cells (S/G2/M phase) among each cluster was quantified. (F) Hierarchy clustering and heatmap of mesenchymal lineage clusters. Color bar on the top indicates the gene expression level. Each cluster contains two batches (top and bottom) of samples.

Using the drop-seq approach, we sequenced the top 1% Td^+^ endosteal bone marrow cells from 1- to 1.5-mo-old *Col2/Td* mice. We profiled 13,759 cells with a median of 2686 genes/cell and a median of 12215 UMIs/cell (Fig. S2A). Unsupervised clustering of the gene expression profiles using Seurat identified 22 groups, including 9 groups of mesenchymal lineage cells and 11 groups of hematopoietic cells, 1 group of endothelial cells, and 1 group of mural cells (Fig. S2B-D). We were surprised to observe a large number of Td^−^ expressing non-mesenchymal lineage cells, which is confirmed by detectable *Td* and *Col2a1* expression in those cells (Fig. S3).

Further analysis of the mesenchymal lineage cells yielded 9 subpopulations (Fig. 1B, total 7585 cells), present in roughly similar proportions across two batches of experiments (Fig. S4). Examination of lineage-specific markers identified clusters with gene signatures of osteoblast (4), osteocyte (5), adipocyte (7), and chondrocyte (8 and 9, Fig. 1C). Identification of chondrocyte clusters, while unexpected, likely represented growth plate chondrocytes released from bone fragments during enzymatic digestion after we cut off epiphyses at the growth plate site. Since chondrocytes originate from the resting zone progenitors but not bone marrow MSCs ^13^, they were excluded from subsequent analysis.

Interestingly, pseudotemporal cell trajectory analysis using Monocle ^14^ placed cells in cluster 1 at one end of pseudotime trajectory and osteocytes (cluster 5) and adipocytes (cluster 7), two terminally differentiated cell types, at the opposite, divergent ends (Fig. 1D). These data suggested that cluster 1 cells are the ancestor of other mesenchymal cells and that they undergo bi-differentiation routes into osteogenic and adipogenic lineage cells. Indeed, cluster 1 expressed several common stem cell markers, such as Sca1, CD34, and Thy1 (Fig. 1C). Compared to cluster 1, clusters 2, 3, and 4 (osteoblast) expressed gradually increased levels of osteogenic genes, such as *Sp7, Alpl, Col1a1, Ibsp*, and *Bglap2*. Since clusters 2 and 3 are sequentially located after cluster 1 and before the branch point in pseudotime trajectory, we named cluster 2 as late MSC and cluster 3 as mesenchymal bi-potent progenitor (MBP). Cells in cluster 6 were distributed around the branch point, indicating that they are lineage committed progenitors (LCPs).

Computational cell cycle analysis was performed to understand the proliferative status of each cluster (Fig. 1E). As a positive control, chondrocyte cluster 9, corresponding to proliferative and prehypertrophic growth plate chondrocytes, had the highest proliferative status. It appeared that within bone marrow mesenchymal lineage cells, MSCs are less proliferative than late MSCs; MBPs, LCPs, and osteoblasts are the most proliferative; adipocytes and osteocytes are non-proliferative. These results further supported our cluster annotation. Hierarchy analysis showed distinct gene expression signatures in each cluster (Fig. 1F).

During the past decade, many proteins have been proposed as MSC markers based on lineage tracing and in vitro culture data, such as LepR ^5^, Mx1 ^15^, Gli1 ^16^, Prxx1 ^17^, Grem1 ^18^, PDGFRα ca1(P S), Nestin, Osterix, CXCL12, and Integrinα /CD200. However, our data suggested that among them, only Sca1 preferentially marks cluster 1. *Pdgfrα, IntegrinαV, Gli1,* and *Prrx1* are broadly expressed across all mesenchymal lineage cells; *Sp7 (Osterix)* and *CD200* are expressed at a higher level in MBPs and LCPs than in MSCs; *Grem1, Lepr,* and *Cxcl12* mark mostly adipocytes and are scarcely expressed in other mesenchymal cells (Fig. S5A); *Nestin* and *Mx1* are mainly expressed in endothelial/mural cells and hematopoietic cells, respectively (Fig. S5B). Hence, the in vivo MSCs we identified here are a stem cell population that are more similar to the previously proposed PαS cells but are different from other proposed MSCs. A previous report demonstrated that sorted PαS cells display mesenchymal progenitor properties in culture and after transplantation ^19^.

### The osteogenic and adipogenic lineage commitment of bone marrow MSCs

Positioning individual cells along a linear pseudotimeline with MSCs as the root revealed transcription factors (TFs) differentially expressed after the branch point of osteogenic and adipogenic lineages (Fig. 2A). Consistent with its longer differentiation pseudotime, adipogenic differentiation required much more unique TFs than osteogenic differentiation, including master regulators *Pparg* and *Cebpα*, several known TFs related to adipogenesis, such as *Klf2, Ebf1,* and *Ebf2* ^24^, as well as many novel ones. Surprisingly, this analysis only revealed master osteogenic regulator *Sp7* but not *Runx2*. Indeed, the expression of *Runx2* started from late MSCs and stayed constant during osteogenic differentiation (Fig. 1C). Our observation that adipogenic differentiation is accompanied with more changes in TFs than osteogenic differentiation is consistent with a recent analysis of in vitro adipogenic and osteogenic differentiation of human MSC-TERT4 cells ^25^.

**Figure 2.**
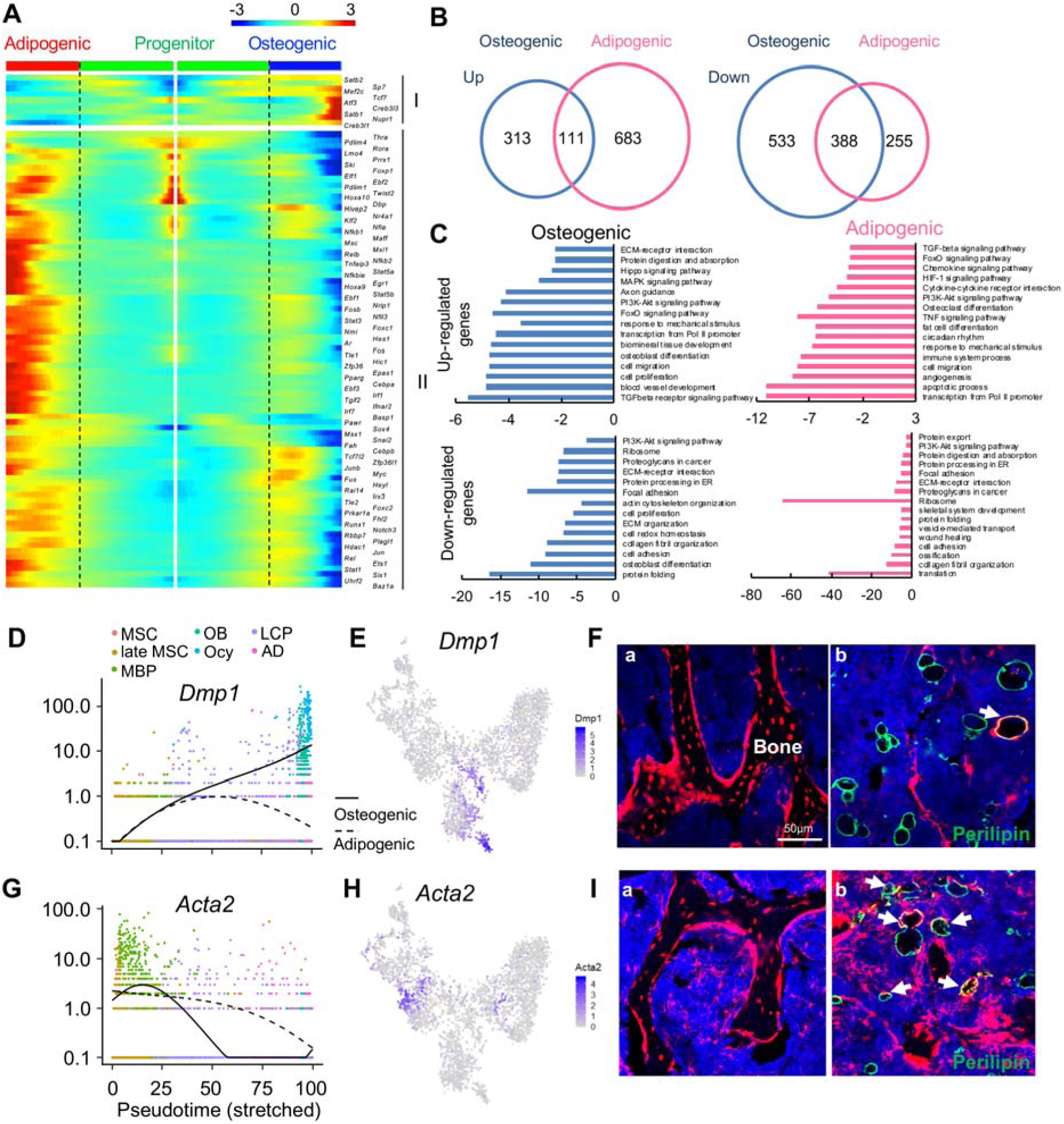
The bifurcated osteo- and adipo-lineage differentiation routes of in vivo bone marrow MSCs. (A) Pseudotemporal depiction of differentially expressed transcription factors (TFs) starting from branching point (dashed lines) toward osteo- (right) and adipo- (left) lineage differentiation. Group I and II contain TFs that are highly up-regulated during osteogenic and adipogenic differentiation routes, respectively. Color bar indicates the gene expression level. (B) Differentially regulated (up-regulated and down-regulated) genes during osteogenic and adipogenic differentiation are counted. (C) GO term and KEGG pathway analyses of genes up-regulated and down-regulated during osteogenic and adipogenic differentiation. Note that some pathways, such as osteoblast differentiation, are identified by both up-regulated and down-regulated genes. This is due to the fact that a set of genes in a pathway are up-regulated while another set of genes in the same pathway are down-regulated. (D) Expression of *Dmp1* goes from the progenitor state and bifurcating into osteogenic or adipogenic branches with respect to pseudotime coordinates. (E) The tSNE plot predicts *Dmp1* expression in osteoblasts, osteocytes, and a portion of LCPs. (F) In 3-month-old *Dmp1/Td* mice, Td labels osteoblasts, osteocytes (a), and only a few adipocytes (b, arrows). (G) Expression of *αSMA* (*Acta2*) goes from the progenitor state and bifurcating into osteogenic or adipogenic branches with respect to pseudotime coordinates. (H) The tSNE plot predicts *αSMA* expression in MBPs. (I) In 4-month-old *αSMAER/Td* mice with Tamoxifen injections at 1 month of age, Td labels osteoblasts, osteocytes (a) and many adipocytes (b, arrows).

Analyzing differentiated expressed genes (DEGs) revealed that up-regulated genes are distinct for each lineage whereas there is considerable overlap between down-regulated genes (Fig. 2B). GO term and KEGG analyses of DEGs revealed unique and common features of osteogenic and adipogenic differentiation processes (Fig. 2C). Some pathways, such as PI3K-Akt, TGFβ, and FoxO signaling pathways, were altered in both lineages, implying their general role in regulating mesenchymal differentiation. Accompanied by downregulation of translation, ribosomal genes were down-regulated in both osteocytes and adipocytes, confirming that they are terminally differentiated, highly specialized cells. The unique features of osteogenic differentiation include extracellular matrix (ECM) organization, ECM-receptor interaction, axon guidance, actin cytoskeleton, biomineral tissue development etc, all of which reflect their bone building function. Interestingly, the unique features of adipogenic differentiation include cytokine-cytokine receptor interaction, immune system process, circadian rhythm, osteoclast differentiation, TNF, HIF-1, chemokines signaling pathways etc. Considering that adipocytes reside in a hematopoietic environment with affluent blood vessels, these features suggest an important regulatory role of adipocytes on its marrow environment.

To validate the lineage differentiation routes predicted by our Seurat and Monocle data, we chose two genes, *Dmp1* and *Acta2 (αSMA)*, for further investigation. DMP1 was previously reported as an osteocyte marker ^26^. Our sequencing analysis revealed that its expression is turned on in LCPs mostly in osteogenic differentiation route (Fig. 2D, E). In 3-month-old *Dmp1-Cre Rosa-tdTomato* (*Dmp1/Td*) mice, Td labeled almost all osteoblasts and osteocytes, some stromal cells but very few Perilipin^+^ adipocytes (6 out of 92 adipocytes counted, n=3 mice, 6.1%, Fig. 2F). αSMA was previously reported as an osteoprogenic marker. We found that it is a marker for the MBP cluster (Fig. 2G, H). In *αSMA-CreER Rosa-tdTomato* (α*SMAER/Td*) mice at 3 months after Tamoxifen injections, Td labeled many osteoblasts, osteocytes (185 out of 256 osteocytes counted, n=3 mice, 68.2%), and bone marrow Perilipin^+^ adipocytes (42 out of 124 adipocytes counted, n=3 mice, 33.7%, Fig. 2I), confirming that αSMA labels mesenchymal progenitors before bifurcated differentiation.

### Age-dependent changes in bone marrow mesenchymal subpopulations

Mouse bone marrow mesenchymal progenitor pool shrinks drastically over the time. Indeed, CFU-F frequency of endosteal bone marrow harvested from 16-month-old *Col2/Td* mice decreased 73% compared to that from 1-month-old mice (Fig. 3A). Similar to adolescent mice, almost all CFU-F colonies from adult and aging mice were Td^+^ (Fig. 3A) and top 1% Td^+^ cells sorted from endosteal bone marrow contained almost all CFU-F colonies (100% Td^+^) in unsorted cells (Fig. S6A, B). These data supported to use a similar scRNA-seq approach on adult and aging bone marrow mesenchymal lineage cells.

**Figure 3.**
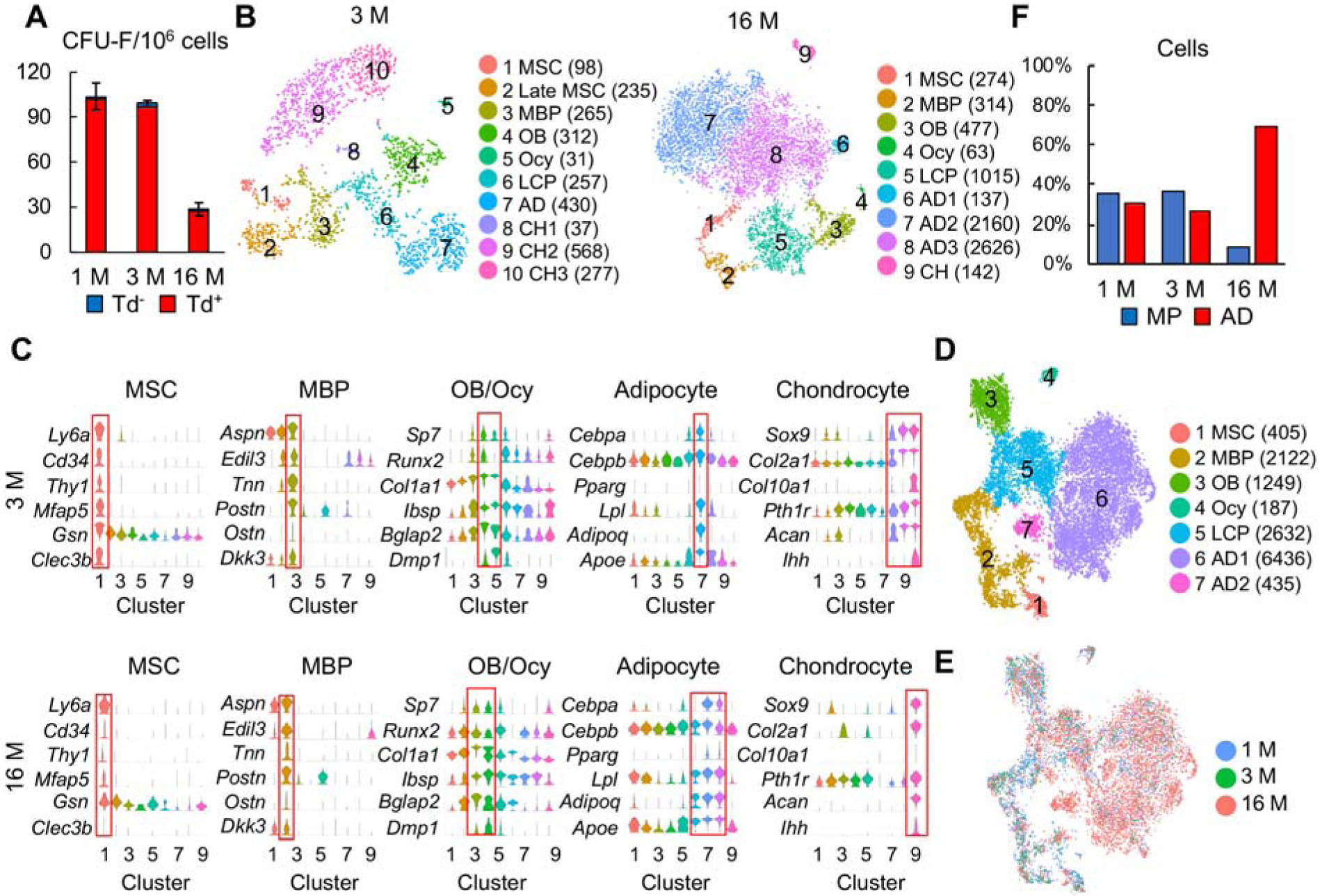
Large scale scRNA-seq analyses of bone marrow mesenchymal lineage cells from 3- and 16-month-old *Col2/Td* mice confirm the same in vivo mesenchymal subpopulations as 1-month-old mice. (A) CFU-F assays of endosteal bone marrow cells from 1-, 3-, and 16-month-old *Col2/Td* mice. n=3 mice/group. Td^+^ and Td^−^ CFU-F colonies are quantified. (B) The tSNE plots of Td^+^ mesenchymal lineage cells isolated from endosteal bone marrow from 3- and 16-month-old *Col2/Td* mice (n=3 mice/group). Cell numbers are listed in parenthesis next to cluster names. MBP: mesenchymal bi-potent progenitor; OB: osteoblast; Ocy: osteocyte; LCP: lineage committed progenitor; AD: adipocyte; CH: chondrocyte. (C) Violin plots of marker gene expression for indicated clusters in 3 and 16 month dataset. (D) An integrated tSNE plot of 1, 3, and 16 month dataset shows the clustering pattern. (E) An integrated tSNE plot of 1, 3, and 16 month dataset shows the distribution of each age group. (F) The percentages of mesenchymal progenitors before lineage commitment (MPs, including MSCs, late MSCs, and MBPs) and adipocytes (ADs) within bone marrow mesenchymal lineage cells are quantified in each age group based on tSNE distribution.

After quality control, we obtained 4502 cells with 19 clusters in 3 month dataset and 8823 cells with 16 clusters in 16 month dataset (Fig. S6C). After removing non-mesenchymal cells, we obtained 2510 (2040 genes/cell and 7243 UMIs/cell) and 7354 (2671 genes/cell and 9660 UMIs/cell) mesenchymal lineage cells in 3 and 16 month dataset, respectively. Unsupervised clustering of cells in both datasets using Seurat yielded similar clustering patterns (Fig. 3B) and cluster markers (Fig. 3C) as 1 month dataset. Notably, late MSC cluster was not detected in 16 month dataset. After removing chondrocytes, pseudotime trajectory again put MSCs at one end and osteocytes and adipocytes at the other two ends in 3 month dataset (Fig. S6D). Interestingly, in 16 month dataset, there was a second branch point separating adipocytes into two ends, most likely due to a drastic increase of adipocyte amount at this age.

Merging all age datasets generated 7 clusters: MSC (1), MBP (1), osteoblast (1), osteocyte (1), LCP (1) and adipocyte (2) (Fig. 3D). It is conceivable to observe that the mesenchymal progenitor pool with bi-lineage differentiation ability was drastically shrunk in the aging sample while the adipocyte population was greatly expanded (Fig. 3E, F), which is consistent with our CFU-F results. Pseudotime trajectory analysis again revealed a similar Y shape curve as 1 month dataset (Fig. S7A). Interestingly, while MSCs and MBPs in 1 and 3 month datasets were more centered at the starting point of pseudotime, those cells in 16 month dataset shifted toward differentiated status, particularly the adipocyte end (Fig. S7B, C). We further examined the expression pattern of adipocyte markers in each cell cluster among different age groups (Fig. S7D). Some of them, such as *Cebpa, Pparg,* and *Lpl,* were expressed at higher levels in most mesenchymal subpopulations, particularly MSC and MBP, in 16 month dataset comparing to 1 and 3 month datasets, suggesting an adipocytic drift during aging. LepR was previously described as a marker labeling adult MSCs but not young MSCs ^5^. As a newly identified marker for the adipocyte cluster, we also observe that its expression is much higher in 16 month MSCs than in 1 month MSCs. Thus, our scRNA-seq data explain why LepR-Cre labels adult MSCs but not young MSCs in mouse bone marrow as described previously ^5^. In summary, we conclude that the same pool of MSCs followed by the same hierarchy differentiation pattern is responsible for bone formation by mesenchymal lineage cells at adolescent, adult, and aging stages. During aging, MSCs are not only reduced in numbers but drifted toward more adipocyte status, which might further account for the loss of progenitor activity.

### Discovery of a novel type of mature adipocyte lineage cells with no lipid accumulation

The conventional view of bone marrow adipocytes is that they are Perilipin^+^ cells with a single large lipid droplet ^28^. However, due to their large size and buoyance, those cells cannot be captured by pelleting and cell sorting. Furthermore, they are rare in the bones of adolescent mice. Therefore, we were surprised to identify a large cell cluster in young mice that express common adipocyte markers, including *Pparg, Cebpa, Adipoq, Apoe,* and *Lpl* (Fig. S8). However, they do not express *Perilipin*, a gene encoding lipid droplet coating protein, and they express *Fabp4,* a gene encoding a fatty acid binding protein, at a very low level, implying that they do not contain lipid droplets. In addition, these cells express a variety of other group defining genes, such as *Lepr, Cxcl12, Il1rn, Serpina3g, Kng1, Kng2, Agt, Esm1,* and *Gdpd2*. qRT-PCR analysis demonstrated that those genes are indeed up-regulated during adipogenic differentiation of mesenchymal progenitors in culture (Fig. S9), supporting that cluster 7 in 1-month dataset represents mature adipocytes.

To validate this population, we constructed mature adipocyte-specific *Adipoq-Cre Rosa-tomato (Adipoq/Td)* mice. In line with the above sequencing data, there were many Td^+^ cells, existing either as stromal cells or pericytes, in the bone marrow of newborn pups (Fig. S10) when long bone undergoes rapid bone formation. At 1 month of age, Td labeled all Perilipin^+^ adipocytes (5 out of 5 cells counted, n=6 mice, 100%), CD45^−^ stromal cells with a reticular shape, and pericytes, but not osteoblasts, osteocytes, periosteal surface, growth plate or articular chondrocytes (Fig. 4A). The majority of Td^+^ cells did not harbor lipid droplets (Fig. 4B, 1995 out of 2000 cells counted, n=6 mice, 99.8%) and none of them incorporated EdU (Fig. 4C, 2060 out of 2060 cells counted, n=3 mice, 0%). After isolated from bone marrow, Td^+^ cells attached to the culture dish but did not form CFU-F colonies (Fig. 4D, 0 out of 100 colonies counted, n=3 mice, 0%). In freshly isolated endosteal bone marrow, Td^+^ cells constituted about 18% of CD45^−^ Ter119^−^ cells. During culture, the percentage of Td^+^ cells decreased progressively after cell passage (P0 when CFU-F colonies became confluent: 2.03%; P1: 0.93%: P2: 0.31%, by flow cytometry analysis), suggesting that those cells have much less proliferative ability if any compared to mesenchymal progenitors. Moreover, while freshly sorted Td^+^ cells from bone marrow of *Col2/Td* mice formed bone with a hematopoietic compartment after transplanted under mouse kidney capsule, freshly sorted Td^+^ cells from *Adipoq/Td* bone did not form bone-like structure (Fig. 4E), demonstrating that they are not mesenchymal progenitors.

**Figure 4.**
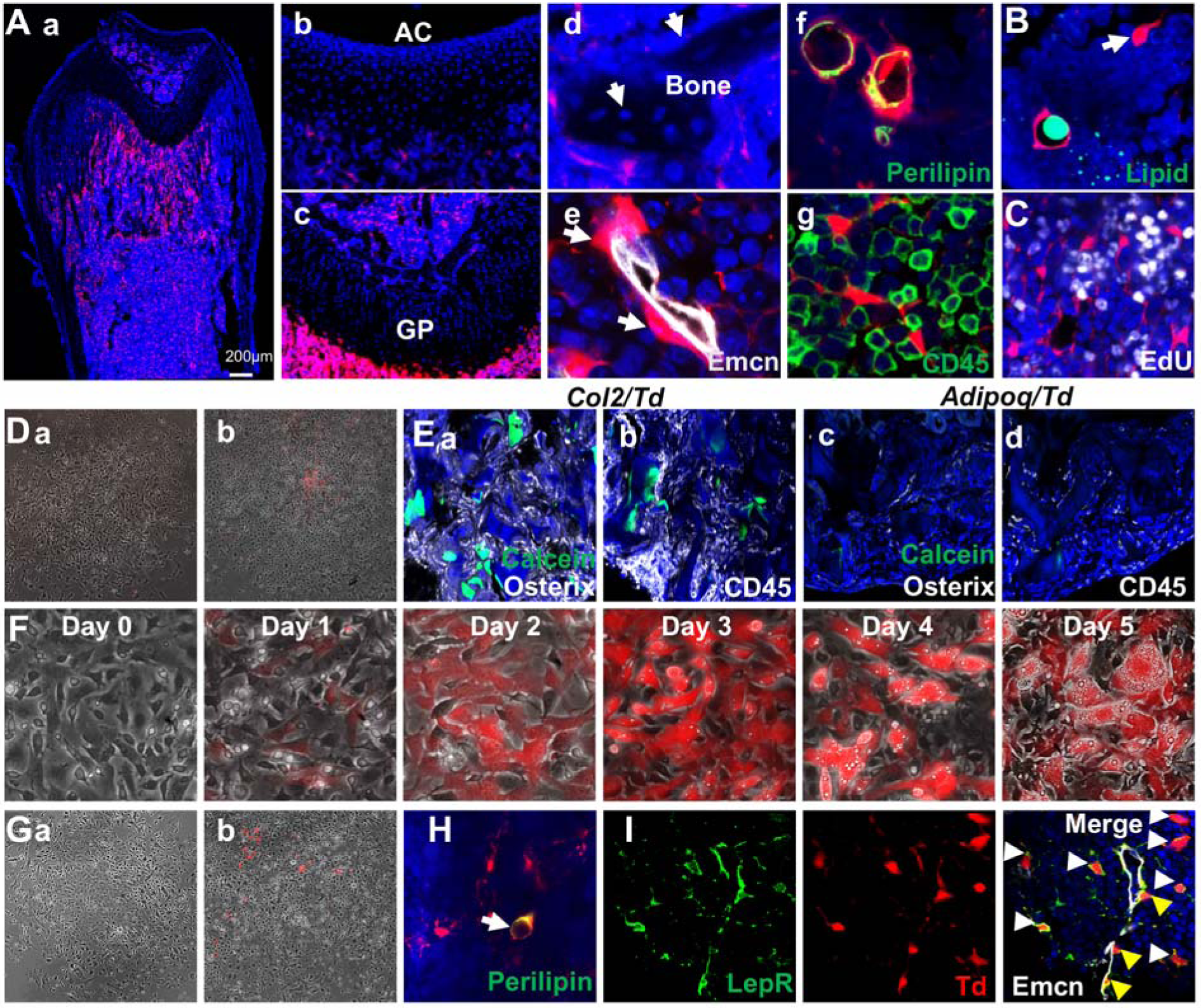
Mouse bone marrow contains abundant non-lipid-laden adipocytes. (A) Representative fluorescent images of 1-month-old *Adipoq/Td* femur reveal many bone marrow Td^+^ cells. (a) A low magnification image of the whole bone. (b-g) At a high magnification, those cells do not label chondrocytes in articular cartilage (b) and growth plate (c), osteoblasts and osteocytes (arrows, d) but label pericytes (arrows, e), Perilipin^+^ adipocytes (f), and CD45^−^ stromal cells (g). (B) According to BODIPY lipid staining, those Td^+^ stromal cells in a reticular shape (arrow) have no lipid accumulation. (C) In vivo EdU injection reveals that Td^+^ cells in *Adipoq/Td* mice do not proliferate. (D) CFU-F assay of bone marrow cells from *Adipoq/Td* mice shows that all CFU-F colonies are made of Td^−^ cells (a). (b) Some Td^+^ cells do attach to the dish but have minimum proliferation ability. n=5 mice. (E) Endosteal bone marrow Td^+^ cells from 1-mo-old *Col2/Td* mice (5 out of 5 transplants), but not *Adipoq-Td* mice (0 out of 3 transplants) form bone-like structure after transplanted under the kidney capsule. Representative fluorescent images of transplants were shown here. Osterix: osteoblasts; CD45: hematopoietic cells; Calcein: new bone surface. (F) In vitro adipogenic differentiation assay demonstrates that non-lipid-laden adipocytes exist as an intermediate state between mesenchymal progenitors and lipid-laden adipocytes. Mesenchymal progenitors, which are Td^−^, were obtained from culturing endosteal bone marrow cells from 1-month-old *Adipoq/Td* mice. Upon confluency, cells were cultured in adipogenic differentiation medium. The same area was imaged daily by inverted fluorescence microcopy. (G) CFU-F assay of bone marrow cells from 1-month-old *AdipoqER/Td* mice (Tamoxifen injections at 2 weeks of age) shows that all CFU-F colonies are made of Td^−^ cells (a). (b) Some Td^+^ cells do attach to the dish but have minimum proliferation ability. n=3 mice. (H) Immunofluorescence staining shows that Perilipin+ bone marrow adipocytes are derived from non-lipid-laden adipocytes in 1-month-old *AdipoqER/Td* mice (Tamoxifen injections at P6, 7). (I) Representative images of co-localization of Td and LepR in stromal (white arrows) and pericytes (yellow arrows) in bone marrow of 1-month-old *Adipoq/Td* mice.

To delineate the relationship among mesenchymal progenitors, non-lipid-laden Perilipin^−^Td^+^ cells, and lipid-laden Perilipin^+^Td^+^ cells, we subjected confluent Td^−^ mesenchymal progenitors from *Adipoq/Td* mice to adipogenic differentiation. Interestingly, Td^−^ cells (day 0) became Td^+^ cells (day 1 and 2) with no lipid droplets first and then evolved into Td^+^ cells with lipid accumulation (Fig. 4F). For fate mapping in vivo, we generated *Adipoq-CreER Rosa-tomato (AdipoqER/Td)* mice. Tamoxifen injections at 2 weeks of age induced many Td^+^ cells in bone marrow at 3 weeks of age with a similar distribution pattern as Td^+^ cells in *Adipoq/Td* mice (Fig. S11). All CFU-Fs from their bone marrow were Td^−^ and only a few Td^+^ cells attached to the dish without further proliferation (Fig. 4G). Since perinatal pups do not have Perilipin^+^ adipocytes in proximal tibiae, Tamoxifen injections at P6-7 label non-lipid-laden adipocytes only. Three weeks later, about 50% of Perilipin^+^ lipid-laden adipocytes in this area were Td^+^ (Fig. 4H), suggesting that non-lipid-laden adipocytes become lipid-laden ones in vivo. Taken together, these in vitro and in vivo data clearly demonstrate that non-lipid-laden adipocytes constitute of a mesenchymal subpopulation situated between mesenchymal progenitors and classic lipid-laden adipocytes along the adipogenic differentiation route of bone marrow MSCs. As a further validation that *Adipoq-Cre* labeled cells are cells in cluster 7 of 1 month dataset, we stained sections with LepR, a cluster 7 marker. As expected, we observed high overlap between Td^+^ cells and LepR^+^ cells in both stromal and pericyte populations (Fig. 4I).

Compared to conventional adipose depots, we did not detect the expression of *Lep* (white adipocyte marker), *Ucp1* (brown adipocyte marker), *Tnfrsf9* (beige adipocyte marker) in cells of adipocyte cluster based on our sequencing data. Other adipose depot markers, such as *Hoxc8, Hoxc9* (white), *Cidea, Cox7a1, Zic1* (brown), *Cited1, Shox2, Tbx1* (beige) ^29, 30^, were expressed either at very low level or ubiquitously among most mesenchymal lineage cells. Overall, our results indicate that bone marrow contains a unique, large population of terminally differentiated adipose-lineage cells that express mature adipocyte markers but do not store significant amounts of lipid.

### A 3D network in bone marrow formed by adipocytic stromal cells and pericytes

In the bone marrow of young *Adipoq/Td* and *AdipoqER/Td* mice, Td^+^ cells exist abundantly as stromal cells and pericytes. PDGFRβ is a general pericyte marker ^31^ and Laminin is a component of basement membrane secreted by both pericytes and endothelial cells ^32^. Our scRNA-seq data showed that both *Pdgfrb* and *Lamb1* are expressed at relatively higher levels in the adipocyte cluster relative to other mesenchymal clusters (Fig. 5A). Pericytes identified by PDGFRβ or Laminin staining in a peri-capillary location were all Td^+^ (850 out of 850 cells counted, n=3 mice, 100%, Fig. 5B), implying that *Adipoq* expression marks all pericytes surrounding bone marrow capillaries. To further confirm their pericyte nature, we mixed sorted Td^+^ cells from *Adipoq/Td* bone marrow with endothelial cells from wildtype bone marrow and conducted vasculogenic tube assays. Strikingly, most Td^+^ cells co-localized with newly formed tubes (Fig. 5C), suggesting that they function as pericytes. Consistent with this, the adipocyte cluster express many angiogenic factors, such as *Vegfa, Vegfc, Angpt4, Rspo3* etc (Fig. 5D). Gene profiling of fractionated Td^+^ cells from *Adipoq/Td* bone marrow confirmed the remarkably high expression of these angiogenic factors as well as adipocyte markers and other newly identified markers (Fig. 5E). The pericyte morphology of adipocytes is unique to bone marrow because we did not observe Td^+^ pericytes in other tissues of *Adipoq/Td* mice, such as muscle, liver, lung, brain, heart, and kidney (Fig. S12).

**Figure 5.**
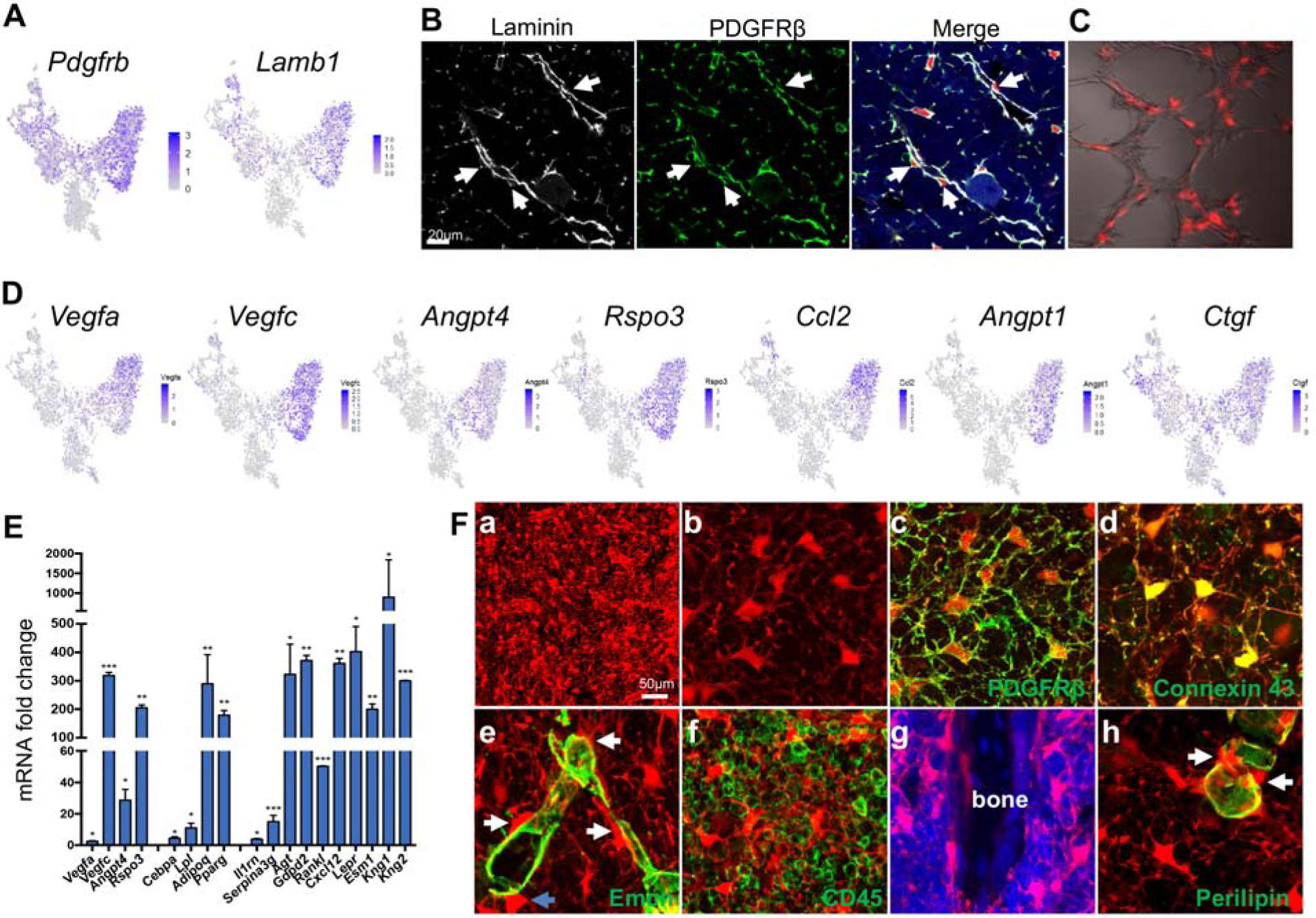
Non-lipid-laden Td^+^ cells in *Adipoq/Td* mice form a vast 3D network through their cell processes ubiquitously distributed inside the bone marrow. (A) The expression patterns of *Pdgfrb* and *Lamb1* are shown in tSNE plots. (B) Immunofluorescence staining reveals that all PDGFRβ^+^ and Laminin^+^ cells with a pericyte morphology are Td^+^ (pointed by arrows). (C) Td^+^ cells sorted from *Adipoq/Td* bone marrow act as pericytes after cocultured with bone marrow endothelial cells undergoing tube formation assay. (D) The expression patterns of adipocyte-secreted angiogenic factors are shown in tSNE plots. (E) qRT-PCR analyses comparing mRNA levels in Td^+^ versus Td^−^ cells (set as 1) sorted from bone marrow of 1-month-old *Adipoq/Td* mice confirm that mature adipocytes highly express angiogenic factors, known adipocyte markers, and novel markers suggested by sequencing data. *: p<0.05; **: p<0.01; ***: p<0.001 Td^+^ vs Td^−^, n=3 mice. (F) A 3D network in bone marrow formed by adipocytic stromal cells and pericytes in 1-month-old *Adipoq/Td* mice. (a) A low magnification image reveals the network of Td^+^ cells in the femoral bone marrow. (b) Td^+^ cell bodies and their processes are more obvious in a high magnification image. See Video1 for a 3D structure. (c) Those Td^+^ cells have PDGFRβ staining all over their cell processes. See Video 2 for a 3D structure. (d) They also have punctuated Connexin 43 staining in their process, indicating cell-cell communication. See Video 3 for a 3D structure. (e) Td^+^ pericytes also have cell processes. Some processes protrude into bone marrow just like those of Td^+^ stromal cells and some of them wrap around the vessel wall. Similarly, Td^+^ stromal cells also extend their processes toward vessels either wrapping around the vessel wall or contacting the processes from pericytes. White arrows point to cells with a typical pericyte morphology. A blue arrow points to a Td^+^ cell sitting on a vessel with a stromal cell shape. Therefore, considering the presence of cell processes and connections, Td^+^ stromal cells and pericytes are indeed very similar. See Video 4 for a 3D structure. (f) Cell processes from adipocytes touch almost every CD45^+^ hematopoietic cells inside bone marrow. (g) They also reach trabecular bone surface. (h) On the contrary, Perilipin^+^Td^+^ adipocytes do not have cell processes. Their cell surface is generally smooth and round. However, we always observed that they are attached to several Td^+^ non-lipid-laden adipocytes (arrows), which have cell processes pointed toward bone marrow. See Video 5 for a 3D structure.

Interestingly, when we scanned 50 µm-thick sections of *Adipoq/Td* long bones, we observed a striking 3D network made of cell processes from non-lipid-laden Td^+^ cells that were ubiquitously distributed inside bone marrow (Fig. 5Fa, b, Video 1). In sharp contrast to round hematopoietic cells, Td^+^ stromal cells and pericytes possessed an average of 6.1±0.2 cell processes (52 cells counted, n=3 mice), which were covered by cell membrane proteins, such as PDGFRβ and gap junction Connexin 43 (Fig. 5Fc, d, Video 2, 3). These processes were reminiscent of osteocytic canaliculi structure, although they are much longer and more disorientated. With these processes, Td^+^ stromal cells and pericytes now are morphologically similar (Fig. 5Fe, Video 4). They all extended cell processes into the marrow, making numerous connections amongst themselves, around endothelial walls (Fig. 5Fe), with hematopoietic cells (Fig. 5Ff), and with bone surface (Fig. 5Fg). Such cell processes were not detected in lipid-laden adipocytes (Fig. 5Fh). The vast 3D structure of Td^+^ cells implies a unique communication role of non-lipid-laden adipocytes.

### The critical function of MERAs in bone

We next sought to study the function of these marrow adipocytes by ablating them in 1-mo-old *Adipoq-Cre Rosa-Tomato DTR* (*Adipoq/Td/DTR*) mice via diphtheria toxin (DT) injections. After two weeks, as expected, the visceral and subcutaneous fat tissues almost disappeared (Fig. S13A). Adipocyte-ablated mice displayed pale bones as compared to mice receiving vehicle injections (Fig. 6A). In bone marrow, both Perilipin^+^ lipid-laden and Perilipin^−^ non-lipid laden Td^+^ adipocytes were greatly reduced in numbers by 95% and 65%, respectively (Fig. 6B, C). Bone marrow vasculature underwent severe pathological changes, becoming dilated and distorted (Fig. 6D, E). Additionally, the number of Emcn^+^CD31^+^ endothelial cells decreased by 45% after DT treatment (Fig. 6F). In the bone marrow, *Adipoq-Cre* labeled a similar density of pericytes as *Col2-Cre* (0.0172±0.0019 vs 0.0174±0.0028 pericytes/µm vessel length, n=3 mice/group). Strikingly, DT injections drastically decreased Td^+^ pericytes by 90% to 0.0018±0.0005 pericytes/µm vessel length (n=3 mice/group, Fig. 6E). Hence, one role of these non-lipid laden adipocytes is to maintain normal bone marrow vessels.

**Figure 6.**
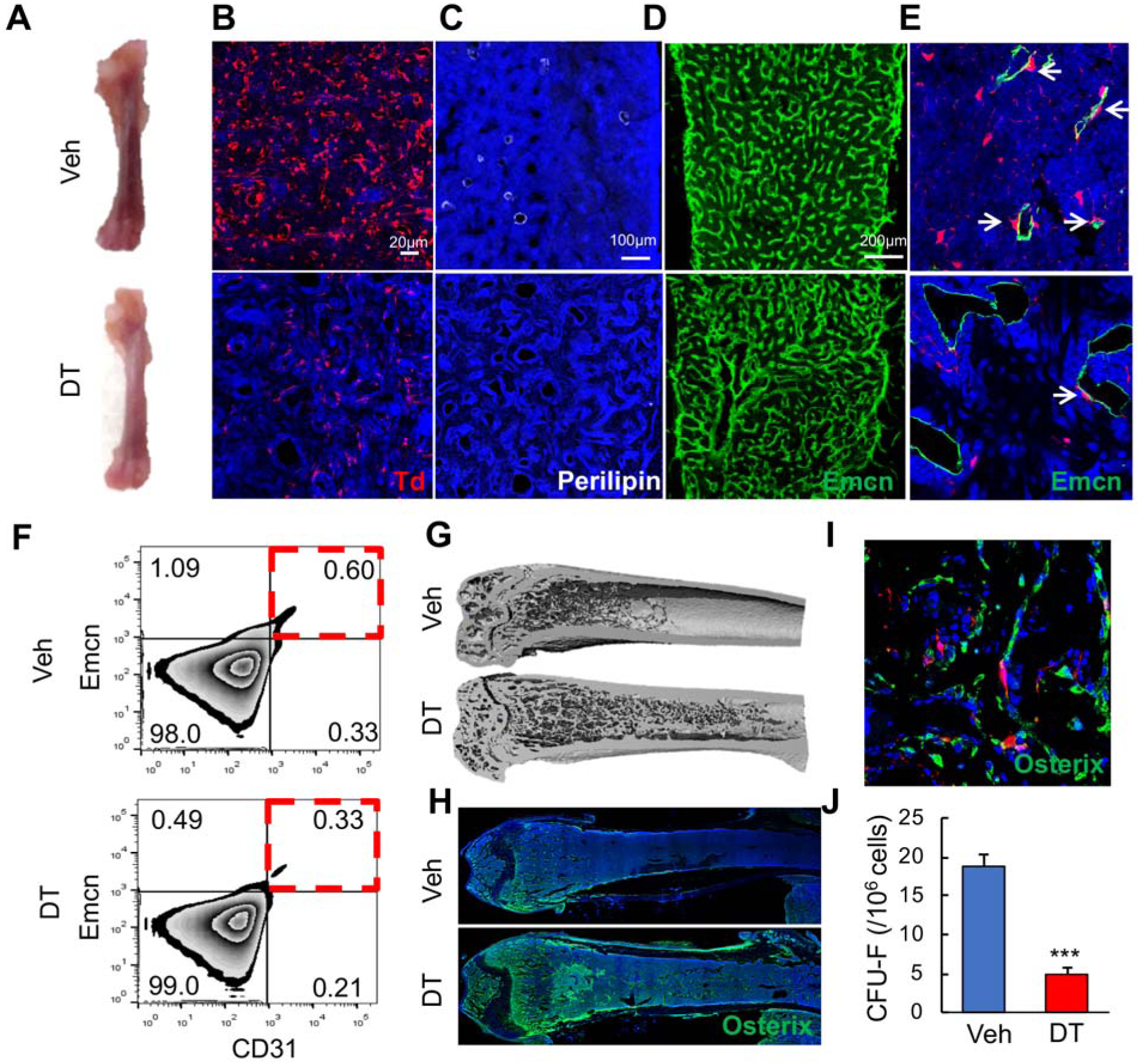
Bone marrow non-lipid-laden adipocytes have important roles in maintaining marrow vasculature and regulating bone homeostasis. (A) Ablation of *Adipoq-Cre* labeled cells changes the color of long bones from red to pink. One-month-old *Adipoq/Td/DTR* mice received either veh or DT injections (50 μg/kg) every other day for 2 weeks and long bones were harvested for analysis. (B) Inside the long bone, Td^+^ cells were significantly reduced by DT injections. (C) Perilipin^+^ adipocytes were also diminished by DT injections. (D) Fluorescent images of vessel staining in femur at a low magnification revealed abnormal bone marrow vessel structure after DT injections. (E) Fluorescent images of vessel staining in femur at a high magnification showed that vessels were dilated coinciding with the depletion of Td^+^ pericytes after DT injections. (F) Flow cytometric analysis of endothelial cells (Emcn^+^CD31^+^) in bone marrow after DT injections. (G) 3D μCT images of *Adipoq/Td/DTR* femurs reveal drastic occurrence of de novo bone formation in the midshaft region after 2 weeks of DT injections. (H) Osterix-stained femoral sections of the entire bones. (I) At a high magnification, fluorescent images clearly show that newly formed osteoblasts (Osterix^+^ cells) in the diaphysis are Td^−^ cells. (J) CFU assay demonstrates a decrease in mesenchymal progenitors in *Adipoq/Td/DTR* femurs after DT injections. n=3 mice/group. ***: p<0.001 DT vs veh.

The color change in bone after adipocyte ablation was due to de novo trabecular bone formation throughout the marrow cavity, especially in the diaphysis that is normally devoid of any trabeculae (Fig. 6G, S13B, C, D). The new bone formation along the endosteal surface of cortical bone increased cortical thickness and area and reduced endosteal perimeter (Fig. S13C, E). Numerous Osterix^+^ osteoblasts were observed inside the diaphyseal bone marrow (Fig. 6H), suggesting that osteogenic differentiation is rapidly activated. The fact that nearly all osteoblasts are Td^−^ (Fig. 6I) further confirmed that *Adipoq-Cre* rarely labels mesenchymal progenitors that generate osteoblasts. Furthermore, DT injections drastically reduced CFU-F frequency from bone marrow (Fig. 6J), suggesting a direct or indirect effect of adipocytes on progenitors. Likely due to the short period, the newly formed bone was mostly woven bone having low tissue mineral density (Fig. S13E) and misaligned collagen fibrils (Fig. S13F). After DT treatment, the overall bone marrow cellularity was significantly decreased (Fig. S14A). However, the hematopoietic components, including hematopoietic stem/progenitors cells and lineage specific cells, as well as peripheral blood components, remained largely unaltered (Fig. S14B, C). Therefore, the reduced cellularity is likely caused by the smaller marrow cavity secondary to the new bone formation.

Non-lipid-laden mature adipocytes were also abundant in the vertebrae of adolescent mice (Fig. S15A). After cell ablation in *Adipoq/Td/DTR* mice, bone marrow vasculature was similarly distorted (Fig. S15B), and osteogenic cells were similarly increased (Fig. S15C) leading to elevated trabecular bone mass (Fig. S15D, E). Note that DT injection alone did not alter bone structure in long bones or vertebrae of *Adipoq/Td* mice.

Taken together, our data indicate that bone marrow non-lipid-laden adipocytes, existing abundantly in both long bones and vertebrae are critical components of the marrow niche. Therefore, we name this new type of adipose-lineage cell as marrow environment regulating adipose cells (MERAs).

### The critical role of MERAs in the bone marrow response to radiation injury

Radiation greatly damages bone marrow and induces marrow adiposity ^11^. To characterize its effects on bone marrow mesenchymal lineage cells, and particularly the newly identified adipocyte population, we focally irradiated the right femurs of 1-month-old *Col2/Td* mice. Unexpectedly, 3 days later, we observed a striking increase of bone marrow Td^+^ cells (non-radiated 1.1±0.1% vs radiated 6.3±0.1%, n=4-6/group, flow cytometry analysis). The similar increases in Perilipin^−^Td^+^ cells (MERAs) was also observed in *Adipoq/Td* mice (Fig. 7A, B). By 7 days post radiation, the number of MERAs returned to a normal level. By contrast, LiLAs steadily increased during this period (Fig. 7A, B), which is consistent with the aforementioned in vitro data that MERAs are precursors of LiLAs (Fig. 4F). Since many MERAs are pericytes, we analyzed Td^+^ pericytes in *Adipoq/Td* mice after radiation. Those cells decreased by 66% along with vessel dilation at 3 days after radiation but returned to normal numbers with relatively normal vessel structure at 7 days after radiation (Fig. 7A, C). While MERAs normally do not proliferate, they started to incorporate EdU shortly after radiation (Fig. 7D). Therefore, the rapid expansion of MERAs after radiation likely contributes to the repair and stabilization of marrow vessels after radiation damage.

**Figure 7.**
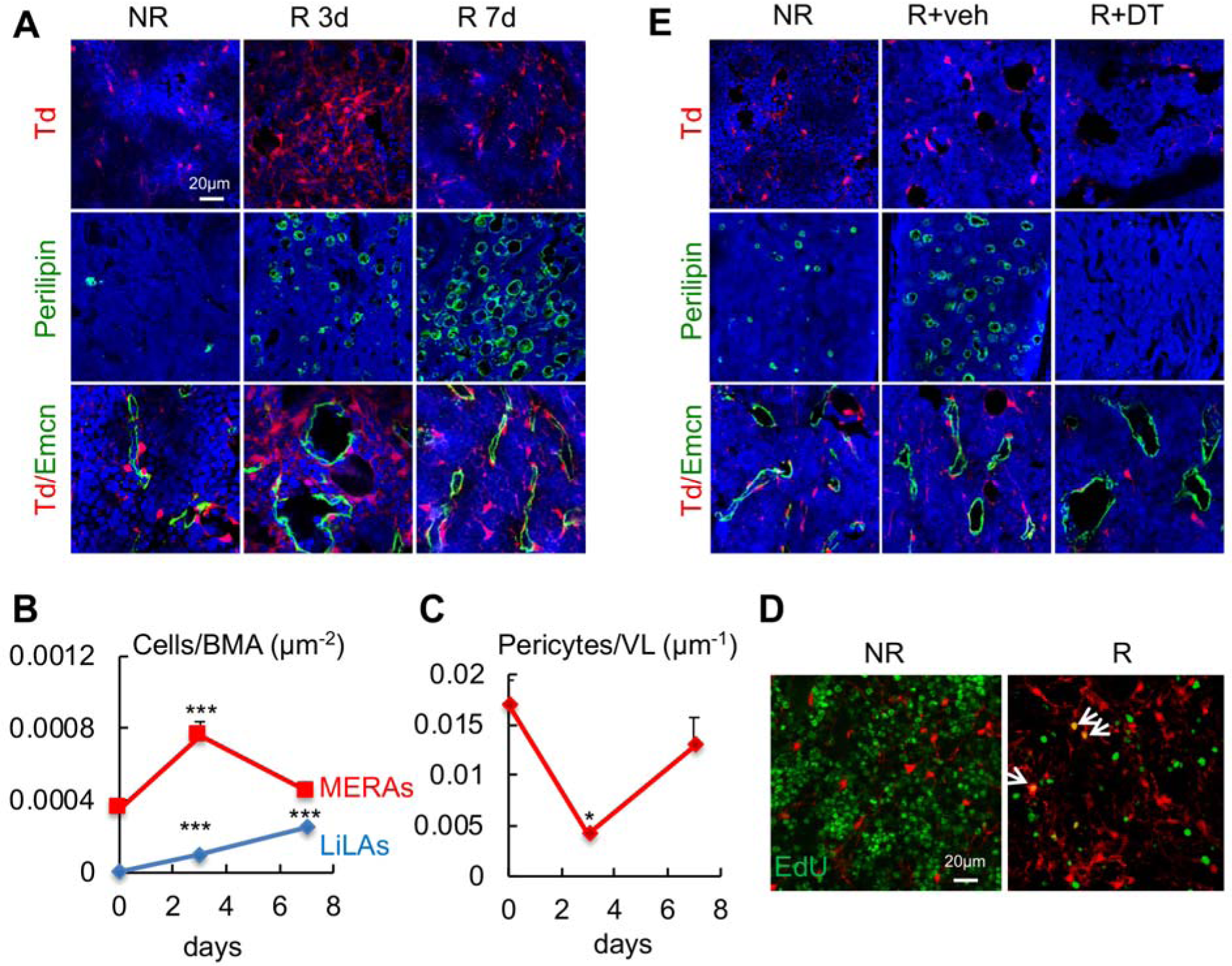
MERAs are required for the recovery of bone marrow vasculature after radiation injury. (A) Representative fluorescent images of Td^+^ cells, Perilipin^+^ LiLAs, and Td^+^ pericytes in the endosteal bone marrow of 1-month-old *Adipoq/Td* femurs before and after focal radiation (3 and 7 days). NR: non-radiation; R: Radiation. (B) MERAs and LiLAs were counted in bone marrow over the time after radiation. BMA: bone marrow area. ***: p<0.001 compared with day 0. (C) MERAs as pericytes were counted in bone marrow over the time after radiation. VL: vessel length. *: p<0.05 compared with day 0. (D) Representative fluorescent images of EdU incorporation in bone marrow cells of *Adipoq/Td* femurs at 3 days post radiation. Arrows point to EdU^+^Td^+^ cells.

Next, we performed large scale scRNA-seq on Td^+^ cells sorted from irradiated endosteal bone marrow of 1-month-old *Col2/Td* mice at 3 days post radiation. We obtained 2014 bone marrow mesenchymal lineage cells. Combining these cells with top 1% Td^+^ mesenchymal cells (2010) from normal 1-month-old mice generated a tSNE plot with a roughly similar cell clusters as before, including MSC, MBP, LCP, adipocytes, osteoblasts, and osteocytes (Fig. S16A). Interestingly, two new clusters, 5 and 6, emerged between MBP and adipocyte clusters. The numbers of radiated cells were significantly lower in MSC and MBP clusters but higher in clusters of 5, 6, and 8 (adipocytes, Fig. S16B). Cell cycle analysis revealed that cluster 6, made of most radiated cells, are highly proliferative (Fig. S16C), which correlates well with the occurrence of EdU^+^ MERA cells detected after radiation (Fig. 7D). Monocle pseudotime analysis suggested that clusters 5 and 6 are differentiated from early and late stages of MBPs, respectively (Fig. S16D). The expression patterns of adipocyte markers, including lipid-associated genes, indicated that both of them are differentiated toward adipocytes (Fig. S16E). Therefore, they were named AP-R1 and R2, respectively. Osteogenic genes, such as Sp7, were expressed more in AP-R2 than R1, indicating that AP-R2 is more differentiated toward osteo-lineage when derived from MBPs.

Taken together, our data revealed the plasticity of mesenchymal differentiation. Radiation damage is able to generate two new adipogenic differentiation routes for MBPs. Among them, AP-R2 is particularly interesting. With an extremely high proliferation, cells in this cluster are essential for generating sufficient amount of MERAs that act as pericytes for repairing damaged vessel structure.

## Discussion

In this study, we computationally delineated the entire in vivo differentiation process of MSCs and defined the mesenchymal lineage hierarchy inside bone marrow (Fig. 8). Stem cell heterogeneity and plasticity have been recognized in some well-studied mammalian tissue stem cells ^33^. However, in the past, owing to a lack of discerning investigative tools, we had ignored these features in mesenchymal lineage cells by simply referring to all progenitors as MSCs or mesenchymal progenitors and searching for one or a set of marker(s) to cover all of them. By identifying the in vivo MSC population and its differentiation route, we have resolved a long standing enigma in bone research that has been plagued by the inconsistency of MSC markers.

**Figure 8.**
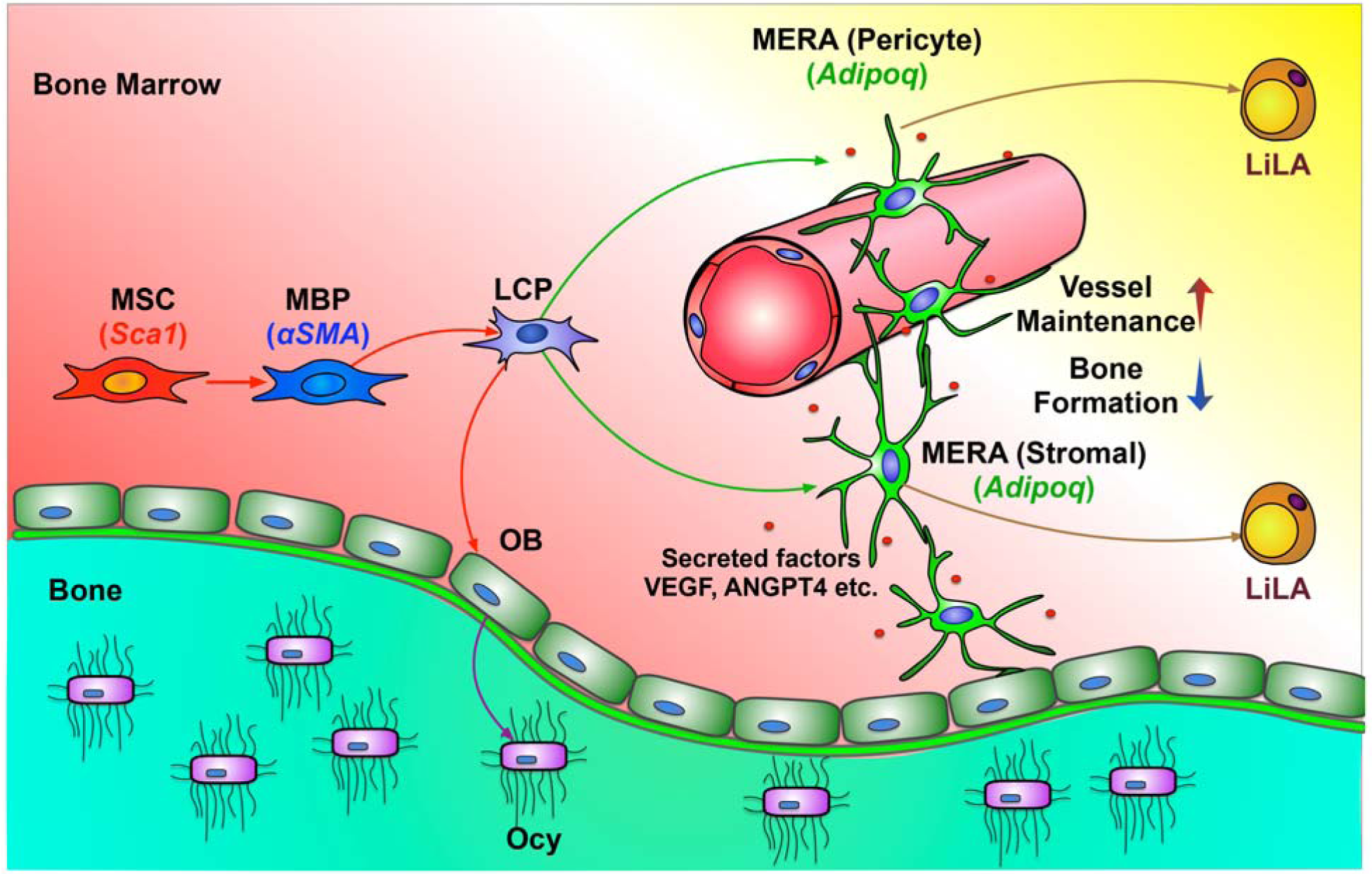
A schematic diagram depicts our model of bone marrow mesenchymal lineage cells and the role of MERAs. MSC: mesenchymal stem cell; MBP: mesenchymal bi-potent progenitor; OB: osteoblast; Ocy: osteocyte; LCP: lineage committed progenitors; MERA: marrow environment regulating adipocytes; LiLA: lipid-laden adipocytes.

It is generally believed that during organ homeostasis, a small number of quiescent stem cells give rise to fast-proliferating progenitor cells, which then mature into various terminally differentiated cell types. This unidirectionality guarantees that different cell types in an organ have distinct identities and play specialized functions. Our proposed in vivo bone marrow MSCs and their bi-lineage differentiation routes are consistent with this stem cell concept. Specifically, this MSC cluster always has the least number of cells among all mesenchymal cell clusters, is always situated at one end of trajectory curve, and shrinks in size during aging. Compared to their progeny progenitors, they are less proliferative and they do not express lineage master regulators, such as *Runx2*, *Sp7*, *Cebpa*, or *Pparg*. Instead, they express constellation of group-defining genes, including *Ly6a*, *CD34*, and *Thy1*, which have been shown to mark other adult stem cells, such as HSCs, keratinocyte stem cells, cancer stem cells, muscle stem cells etc ^34–37^. Along the differentiation routes of MSCs, the expression levels of lineage specific genes, such as *Bglap2*, *Col1a1*, *Ibsp*, *Adipoq,* and *Lpl*, progressively increase, confirming an undifferentiated nature of MSCs. Our definition of MSCs is also consistent with a previous study showing that mouse bone marrow Sca1^+^PDGFRα^+^CD45^−^Ter119^−^ cells generate colonies at a high frequency and differentiate into hematopoietic niche cells, osteoblasts, and adipocytes after in vivo transplantation ^19^.

A recent report examined the same question by performing scRNA-seq on Ter119^−^CD71^−^Lin^−^ bone marrow cells from *WT* mice ^8^ and generated a similar clustering pattern and gene expression signature as ours, demonstrating the robustness of scRNA-seq methodology of identifying bone marrow mesenchymal populations. Based on a previous study showing that LepR labels adult bone marrow MSC ^5^, a Lepr^+^ cell cluster was assigned as MSCs. However, this cluster is the largest one among all mesenchymal lineage clusters, which is unusual for a stem cell cluster. Moreover, it also highly and specifically expresses many markers of MERAs, including known adipocyte markers *Adipoq, Pparg, Apoe, Cebpa, Cebpb, Lpl* (Fig. S17A), suggesting that they represent adipose cells. Interestingly, among 5 fibroblast cell clusters identified in that study, 2 small ones have a similar set of gene signatures as our MSCs and the rest 3 ones overlap with MBPs. One osteolineage cluster in their dataset shares similarity to our LCPs and the other one is similar to our osteoblasts (Fig. S17B).

Using a lineage tracing approach (*Lepr-Cre* mice), Zhou et al.^5^ previously showed that LepR labels bone marrow mesenchymal progenitors in adult mice (6 months of age) but not in young mice. Our study identified Lepr is a marker for MERAs, which are non-proliferative and express mature adipocyte genes. We also observed that *Lepr* express scarcely in 1 and 3 month MSC clusters and highly in 16 month MSCs (Fig. S7D, S8B). The increased *Lepr* expression in our MSCs during aging might explain the previous observation that LepR marks adult MSCs but not young MSCs. In the same study, it was reported that LepR^+^ cells constitute 0.3% of bone marrow cells. Since bone marrow CFU-F frequency is only 0.001-0.01% in adult mice, these data clearly indicate that LepR labels much more than progenitors. Our aging sequencing data showing the great expansion of LepR^+^ MERA cluster in 16-month-old mice further excludes the use of LepR as an MSC marker because it conflicts with the well-known aging effect on reducing the pool of bone marrow mesenchymal progenitors. Therefore, we believe that LepR^+^ cells in bone marrow are a heterogeneous population whose majority is MERAs and minority is mesenchymal progenitors. This is further confirmed by another scRNA-seq study analyzing *Lepr-Cre* labeled bone marrow cells ^7^. In that study, two large clusters, Mgp^High^ P1 and Lpl^High^ P2, were found to highly express adipogenesis-associated genes (Fig. S18). The other two small clusters are similar to our LCP and OB clusters.

Bone marrow adipocytes are conventionally viewed as large cells containing a unilocular lipid droplet and their functions in bone metabolism are debated ^28^. Based on our computational data, we uncovered a novel type of adipose-lineage cell, MERAs, that expresses many markers of terminally differentiated adipocytes, but lack significant lipid stores. Our trajectory analysis, in vitro adipogenic differentiation assay, and in vivo fate mapping data provide strong evidence that MERAs represent a stable transitional cell type along the adipose differentiate route after mesenchymal progenitors and before LiLAs. In traditional adipose depots, adipogenic differentiation is tightly coupled with lipid accumulation. Single cell analyses of murine or human white fat depots (visceral or subcutaneous) did not identify cell populations analogous to MERAs ^38^. Pre-adipocytes from adult tissues express low levels of *Pparg* but not *Adipoq*, and are also defined by their expression of canonical mesenchymal progenitor markers, including *Sca1* and *Cd34* ^39^. In contrast, MERAs do not proliferate and they express many markers of terminally differentiated adipocytes. The *Cre*s that label MERAs, *Adipoq-Cre* and *CreER,* are highly specific for mature adipocytes in other adipose depots because they do not label stromal vascular fraction (SVF) containing pre-adipocytes ^40^. Therefore, MERAs represent a distinct cell type from traditional pre-adipocytes. Under physiology conditions, MERAs are much more abundant than LiLAs (400-fold in 1-month-old mice), indicating that they are a stable population rarely converted into LiLAs. Whether they can directly undergo apoptosis without differentiated into LiLAs needs further investigation. It will also be interesting to identify the molecular mechanism that blocks so many adipose lineage cells at MERA stage without becoming LiLAs. We speculate that MERAs are poised to accumulate lipid and convert to LiLAs in response to metabolic or other cues, such as radiation injury and the availability of fatty acids. Overall, we conclude that MERAs represent a marrow-specific type of adipose cell that plays unique roles in regulating their bone environment.

Morphologically and functionally, MERAs are also different from traditional round shape adipocytes. Even in young mice, MERAs exist in abundant amount as marrow stromal cells or pericytes. Importantly, their cell processes form a vast 3D network structure inside bone marrow, making numerous contacts among themselves and with the rest of bone marrow components. Most likely through secreting factors into their marrow environment, they play pivotal roles in maintaining marrow vasculature, suppressing osteogenic differentiation of mesenchymal progenitors. Although we did not detect any changes in hematopoietic components shortly after ablation of MERAs, we still believe that they have the ability of controlling hematopoiesis because of several reasons. First, Cxcl12, a chemokine responsible for the retention of hematopoietic progenitors ^41^, is the 2^nd^ most expressed genes in MERAs. Previous studies have shown that CAR cells act as a niche for HSCs ^42^. Second, it has been shown that Scf from *Adipoq-CreER* or *Lepr-Cre* labeled bone marrow cells promotes hematopoietic regeneration after injury ^43^. Indeed, according to our datasets, Scf is highly and specifically expressed in MERAs. Hence, a prolonged cell ablation period or an injury might be required to detect the regulatory action of MERAs on hematopoiesis inside the bone marrow.

In the literature, pericytes refer to microvascular periendothelial cells that are embedded within the vascular basement membrane. Currently, the relationship between pericytes and MSCs in bone marrow is still ambiguous. Since LepR^+^ cells and CAR cells marks bone marrow cells and some of those cells are perivascular, it was proposed that pericytes are a heterogenic population containing MSCs. However, similar to MERAs, LepR^+^ cells and CAR cells also exist as stromal cells. Without a specific marker to distinguish between pericytic and stromal LepR^+^/CAR cells, it is difficult to point out where MSCs are located. Using *Adipoq-Cre(ER)* reporter system, we are able to conclude that pericytes do not contain MSCs based on the following reasons. First, staining of multiple pericyte markers PDGFRβ and basement membrane component Laminin revealed that all pericytes surrounding Emcn^+^ capillaries are Td^+^ cells in *Adipoq/Td* mice. These data point out that bone marrow pericytes are a relatively homogenous population made of non-lipid laden adipocytes. In our previous study of radiation damage on bone, we noticed that some radiation-induced LiLAs make direct contacts with blood vessels in such a way that they look like budding from the vessels ^11^. With our current data, it is reasonable to assume that radiation promotes the differentiation of pericytic MERAs into LiLAs. Second, our data revealed that MERAs are not proliferative. We are aware that our *AdipoqER/Td* data conflict with a previous study suggesting that *Adipoq-CreER* labeled cells are adipoprogenitors with CFU-F forming ability ^43^. The discrepancy might lie in different ages of mice, different bone marrow harvest methods, different culture conditions, and probably different criteria of defining CFU-Fs. Nevertheless, our CFU-F data match with our computational cell cycle results, in vivo EdU incorporation, and the general consensus that *Adipoq-Cre(ER)* labels mature adipocytes only ^40^. Compared to other Cres, we believe that *Adipoq-Cre* provides a more defined system to label and characterize pericytes.

Our study of *Adipoq(ER)/Td* mice challenges another pericyte concept that pericytes must be tightly associated with endothelial cells and embedded within the basement membrane synthesized by both pericytes and endothelial cells. Reviewing the 3D images of bone marrow from *Adipoq/Td* mice clearly indicated that some perivascular Td^+^ cells do not have the authentic pericyte morphology. Instead, they have stromal cell morphology but contact endothelial cells through their cell bodies and processes. Moreover, many Td^+^ stromal cells having no cell body contact but extend their cell processes to wrap around capillaries walls, presumably also contributing to vessel stabilization and maintenance. Therefore, the concept of pericytes based on its location and morphology should be revisited to suit the special feature of MERAs in bone.

One limitation of our study is that we do not have a Cre specific for non-lipid-laden adipocytes inside bone marrow. Therefore, we could not rule out the potential influence of lipid-laden adipocytes (LiLAs). Since LiLAs do not have cell processes (Fig. 5Hh, Video 5) and MERAs are much more abundant than LiLAs at 1 month of age, we believe that the later one has a minor role if any.

The main functions of currently known adipocytes are lipid storage, thermogenesis, and regulation of nutrient homeostasis. Apparently, compared to them, MERAs possess a completely different set of functions. Why bone marrow mesenchymal progenitors need to turn on adipocyte transcriptome in order to become a central regulator of bone marrow environment is an interesting question. Furthermore, future studying the underlying molecular mechanisms by which they regulate their environment will shed light on identifying new drugs for treating bone diseases such as osteoporosis.

## Acknowledgments

We thank Dr. Henry Kronenberg from Harvard University for critics and guidance on this study. We also thank Dr. Ivo Kalajezc from University of Connecticut Health Center for giving us α*SMA/Tomato* mice.

## Funding

This study was supported by NIH grants NIH/NIAMS R01AR066098, R01DK095803, R21AR074570 (to L.Q.), P30AR069619 (to Penn Center for Musculoskeletal Disorders), American Heart Association 17GRNT33650029 (to Y.G.), R01HL095675, R01HL133828, DOD (to W.T.), NRSA F31HL139091 (to N.H.), NIH/NIDCR R00DE025915, R03DE028026 (to C.C.).

## Author contributions

L.Z., L.Y., R.J.T., and L.Q. designed the study. L.Z., L.Y., Y.W., C.C., W.Y., R.J.T., and L.W. performed animal experiments. R.S. helped with library construction. L.Z., L.Y., Y.W., and W.Y. performed histology and imaging analysis with assistance from R.J.T. L.Z., L.Y., and N.H. performed FACS experiments. L.Z., L.Y., L.W., and W.Y. performed cell culture and qRT-PCR experiments. R.J.T., L.Y., and L.Q. performed computational analyses with assistance from Z.M., J.P, and M.L. Y.Z. W.T., Y.G., J.A., F.L., P.S., K.S., M.L., and C.C. provided administrative, technical, or material support and consultation. L.Q. wrote the manuscript. L.Z., L.Y., J.A., F.L. and P.S. reviewed and revised the manuscript. L.Q. approved the final version.

## Competing interests

Authors declare no competing interests.

## Materials & Correspondence

should be addressed to L.Q.

## Data and materials availability

All data is available in the main text or the supplementary materials.

## Extended data

Materials and Methods

Figures. S1 to S18

Tables S1

Videos: 1-5

References (1-18)

## Notes

#### Summary of Updates

Figure S11 legend revised.

