## supplementary material for "Single cell transcriptomics identifies a unique adipocyte population that regulates bone marrow environment"

**This PDF file includes:**

Materials and Methods

Figs. S1 to S18

Tables S1

Materials and Methods

*Animal study design*

All animal work performed in this report was approved by the Institutional Animal Care and Use Committee (IACUC) at the University of Pennsylvania. In accordance with the standards for animal housing, mice were group housed at 23-25°C with a 12 h light/dark cycle and allowed free access to water and standard laboratory pellets.

*Col2-Cre Rosa-tdTomato* (*Col2/Td*), *Dmp1-Cre Rosa-tdTomato* (*Dmp1/Td*), *Adipoq-Cre Rosa-tdTomato* (*Adipoq/Td*), *Adipoq-CreER-tdTomato (AdipoqER/Td)* mice were generated by breeding *Rosa-tdTomato* (Jackson Laboratory, Bar Harbor, ME, USA) mice with *Col2-Cre* ^1^ (Jackson Laboratory), *Dmp1-Cre* ^2^ (Jackson Laboratory), and *Adipoq-Cre* ^3^ (Jackson Laboratory) mice, *Adipoq-CreER* ^4^ (Jackson Laboratory) respectively. *αSMA-CreER Rosa-tdTomato* (*αSMAER/Td*) was a gift from Dr. Ivo Kalajezc ^5^. For lineage tracing experiments, *αSMAER/Td* mice received Tamoxifen injections (75 mg/kg/day) at 1 month of age for 2 days and their bones were harvested at 3 months later. For CFU-F experiments, *AdipoqER/Td* mice received tamoxifen injections (75 mg/kg/day) at 2 weeks of age for 3 days and their bones were harvested 1 week later. For lineage tracing experiment, *AdipoqER/Td* mice received Tamoxifen injections (75 mg/kg/day) at P6 and P7 and their bones were harvested at 1 month of age. *Adipoq-Cre Rosa-tdTomato DTR* (*Adipoq/Td/DTR*) mice were generated by breeding *Adipoq/Td* mice with *Rosa-DTR* mice (Jackson Laboratory). Mice received vehicle (1xPBS) or DT injections (50 µg/kg) every other day for 2 weeks. Bones were harvested at indicated times for histology and microCT analysis.

For focal radiation, mouse right femur received a clinically relevant radiation dose of 5 Gy using small animal radiation research platform (SARRP, Xstrahl, Suwanee, GA) as we described previously ^6^. The radiation was delivered to the distal femur (15 mm in diameter) at a rate of 1.65 Gy/min with the aid of built-in µCT and X-ray. For cell ablation experiment, DT injections started right after radiation.

For in vivo transplantation, freshly FACS-sorted Td^+^ cells (5x10^4^/transplant) were mixed with Gelfoam and placed under the kidney capsule of recipient 2-month-old *C57Bl/6* mice. The transplant grafts were harvested 4 weeks post-transplantation for histology analysis. Mice received calcein injection (15 mg/kg) at 1 day prior to euthanization.

*Endosteal bone marrow Td^+^ cell isolation and cell sorting*

Endosteal bone marrow cells were harvested as described previously ^7^. Briefly, the outer surfaces of long bones were scraped and digested to remove the periosteum. After cutting off the epiphyses and flushing out the central bone marrow, metaphyseal bone fragments were longitudinally cut into two halves and digested by proteases to collect endosteal bone marrow cells. Freshly isolated endosteal bone marrow cells were resuspended into FACS buffer containing 25 mM HEPES (Thermofisher scientific) and 2% FBS in PBS and sorted for top 1% Td^+^ cells if a Td peak was not obvious or Td^+^ cells if a Td peak was obvious using Influx B (BD Biosciences, San Jose, CA) or Aria B (BD Biosciences, San Jose, CA).

*Single-cell RNA sequencing of endosteal bone marrow cells*

We constructed 5 batches of single cell libraries for sequencing: endosteal Td^+^ bone marrow cells from 1-month-old (n=2), 1.5-month-old (n=3), 3-month-old (n=3), 16-month-old (n=3) and 1-mon-old irradiated (n=3) male *Col2/Td* mice. Libraries were generated by Chromium controller (10X Genomics Inc, San Francisco, USA), barcoded and purified as described by the manufacturer, and sequenced using a 2x150 pair-end configuration on an Illumina HiSeq platform at a sequencing depth of ~400 million reads. Cell ranger was used to demultiplex reads, followed by extraction of cell barcode and unique molecular identifiers (UMIs). Read 2 was aligned to a modified reference mouse genome (mm10) using STAR ^8^. Doublets or cells with poor quality (genes>6000, genes<200, or >5% genes mapping to mitochondrial genome) were excluded. Expression was natural log transformed and normalized for scaling the sequencing depth to a total of 1x10^4^ molecules per cell, followed by regressing out the number of UMIs and percent mitochondrial genes using Seurat ^9^.

For the integrated dataset, canonical component analysis (CCA) ^10^ was performed using the union of the top 2,000 genes with the highest dispersion from both datasets and to determine the common sources of variation between 1- and 1.5-month datasets and between 1-month normal and irradiated datasets. A CCA dimensional reduction was subsequently generated on the basis of the first 30 canonical correlation vectors. T-distributed stochastic neighbor embedding (tSNE) plots were used to visualize the data based on the CCA alignment. Cluster specific markers for each cluster, relative to the remaining population, were conducted using the bimodal analysis to identify differentially expressed genes (DEGs). Sub-clustering was performed by isolating the mesenchymal lineage clusters identified from the remaining bone marrow cells using known marker genes, followed by reanalysis as described above. This resulted in the generation of 9 clusters. Because chondrocytes are likely derived from the growth plate, not endosteal MSCs, these chondrocyte clusters were excluded from further analyses. DEGs between these clusters were generated as described above. GO terms and clusters, as well as KEGG pathway enrichment, were identified using the database for annotation, visualization and integrated discovery (DAVID) ^11^.

For individual analysis of the 3- and 16-month-old dataset, Seurat package was used for filtering, variable gene selection, dimensionality reduction analysis and clustering standardly. Poor quality cells or doublets were first filtered as described above, followed by identifying the variable genes by controlling for the relationship between average expression and dispersion. Next, we performed principal component analysis (PCA) using the JackStraw function. Statistically significant PCs were selected as input for tSNE plots. For sub-clustering, we repeated the same procedure of ﬁnding variable genes, dimensionality reduction, and clustering. Different resolutions for clustering have been tested to demonstrate the robustness of clusters.

To analyze 4 datasets from different age group, we applied Seurat V3^12^ for performing integrated analyses to identify common cell types. Anchors from different dataset were defined using the FindIntegrationAnchors function, then use these anchors to integrate the 4 datasets together with IntegrateData. This integrated data was further analyzed as described above.

For cell cycle analysis, we used a core set of 43 S and 54 G2/M genes defined previously ^13^. First, the genes that are expressed in less than 5% of total cells were removed, resulting in 34 S and 45 G2/M genes for the following cell cycle analysis. Second, we define proliferative cells if the cell express S gene set or G2/M gene set, that is, there is a significant difference between the expression value of S genes and G2/M genes. Since the null distribution for the S gene set and G2/M gene set is unknown, we designed a permutation test which does not assume a null-distribution. To be more specific, for each cell, we resampled 34 genes from the 79 genes (34+45) as "S genes" and the rest 45 genes as "G2/M genes". This resampling was repeated 5000 times and each time, we calculate the difference between the mean expression value of "S genes" and the mean expression values of "G2/M genes". The original difference between the mean expression value of S genes and the mean expression value of G2M was compared with the difference between resampling results to get a significant score (p-values). To correct for multi-test, FDR corrections were applied to each cluster. At last, FDR p-value < 0.05 was used as a cutoff for distinguishing proliferative or non-proliferative cells.

To computationally delineate the developmental progression of bone marrow mesenchymal cells and order them in pseudotime, we used the algorithms implemented in the Monocle package ^14^. We include the mesenchymal lineage cells with no chondrocytes from separated or integrated dataset of different age (1, 3 and 16) month old or 1 month old normal/irradiated mice for the analysis. We ordered cells by selecting genes with high dispersion across cells, using a parameter of “mean_expression >= 0.05 & dispersion_empirical >= 2 * dispersion_fit”, lists of genes were selected for dimensional reduction to generate the trajectory reconstruction using the nonlinear reconstruction algorithm DDRTree. Branched expression analysis modeling, or BEAM ^14^ was used to determine the genes that are differentially expressed between the osteogenic or adipogenic branches. A mouse transcript factors (TF) list (TFdb) ^15^ were used to detect 385 TFs among the differential expressed genes.

*Histology*

To obtain whole mount sections for immunofluorescent imaging, freshly dissected bones (femurs, tibiae, L4/L5 vertebrae) were fixed in 4% PFA for 1 day, decalcified in 10% EDTA for 4-5 days, and then immersed into 20% sucrose and 2% polyvinylpyrrolidone (PVP) at 4°C overnight. Then sample was embedded into 8% gelatin in 20% sucrose and 2% PVP embedding medium. Samples were sectioned at 50 µm in thickness. Sections were incubated with goat anti-LepR (R&D system, AF497), rabbit anti-Laminin (Sigma, L9393), rat anti-CD45 (Biolegend, 103101), rat anti-Endomucin (Santa cruz, sc-65495), rabbit anti-Osterix (Abcam, ab22552), rabbit anti-Perilipin (Cell signaling, 9349), rat anti-mouse PDGFRβ (Biolegend, 136002), rabbit anti-mouse connexin 43 (Cell signaling, 3512) at 4°C overnight followed by Alexa Fluor 488-conjugated donkey anti-goat (Abcam, ab150129), Alexa Fluor 647 anti-rat (Abcam, ab150155) or anti-rabbit (Abcam, ab150157) secondary antibodies incubation 1 hour at RT. For lipid staining, BODIPY dye (ThermoFisher scientific, D3822) was added together with secondary antibody and incubate for 1 hour at RT. For EdU staining, mice received 1.6 mg/kg EdU 1 day and 3 hr before sacrifice and the staining was carried out according to the manufacturer’s instructions (ThermoFisher scientific, Click-iT™ EdU Alexa Fluor™ 647 Imaging Kit, D3822).

To reconstruct 3D structure of adipocytes, fluorescence images were captured by a Zeiss LSM 710 scanning confocal microscope interfaced with the Zen 2012 software (Carl Zeiss Microimaging LLC, Thornwood, NY). Confocal image stacks were collected to a depth of ~50 µm and a step size of 1 µm at 63x magnification. Laser power and detector sensitivity were adjusted for z-correction to compensate for signal dissipation at greater imaging depths. Detector gain and offset were adjusted according to the most intense regions to ensure minimal saturation of the signal over the entire imaging area. Imaris 9.2 software (Oxford instruments, Switzerland) was used for processing z-stack and generating movies. The number of cell processes per cell was manually quantified.

*Cell culture*

For CFU-F assay, unsorted endosteal bone marrow cells were plated at 1x10^6^ cells/T25 flask. Sorted endosteal bone marrow cells were plated at 1x10^4^, 1x10^4^, and 9.8x10^5^ cells/T25 flask for top 1%, 1-2%, and >2% group, respectively, based on Td intensity. Cells were cultured in growth medium (α-MEM supplemented with 15% FBS, 0.1% β-mercaptoethanol, 20 mM glutamine, 100 IU/ml penicillin, and 100 µg/ml) for 7 days before counting CFU-F number.

*Tube formation assay*

16-well slide chamber was coated with 60 μl matrigel (Corning, 356231) per well on ice, and polymerize at 37°C for 30 min. Endothelial Progenitor Outgrowth Cells (EPOC) were cultured in growth medium (mouse EPOC basal medium with 10% FBS, 100 IU/ml penicillin, and 100 µg/ml streptomycin). Once confluency, mix 350000 EPOC cells with 14000 freshly sorted Adipoq/Td^+^ cells, seeded mixed cells into matrigel coated slide chamber. After 8 hours, cells were observed under fluorescence microscopy.

Mesenchymal progenitors were obtained by culturing endosteal bone marrow cells at a high density (3x10^6^ cells/T25 flask). Once confluent, cells were cultured in adipogenic medium (DMEM with 10% FBS, 10 ng/ml triiodothyronine, 1 µM rosiglitazone, 1 µM dexamethasone, 10 µg/ml insulin, 100 IU/ml penicillin, and 100 µg/ml streptomycin) for 7 days. Brightfield and fluorescent images of mesenchymal progenitors from 1-month-old *Adipoq/Td* mice undergoing adipogenic differentiation were taken from day 0 to 5 by fluorescence inverted microscopy (Nikon Eclipse, TE2000-U).

*Flow cytometry*

Freshly isolated endosteal bone marrow cells were centrifuged to pellet cells. ACK lysing buffer (ThermoFisher Scientific, A1049201) was added to lyse red blood cells and then stained with Endomucin (FITC rat anti-mouse Endomucin, Santa cruz biotechology sc-65495) and CD31 (APC rat anti-mouse CD31, Biolegend, 561814) antibodies for 45 min on ice. Wash with flow buffer (2% FBS in PBS) twice. Sample was run on LSR A. Data was analyzed by FlowJo X.

*qRT-PCR analysis*

Sorted cells or cultured cells were collected in TRIzol Reagent (Sigma, St. Louis, MO, USA). A Taqman Reverse Transcription Kit (Applied BioSystems, Inc., Foster City, CA, USA) was used to reverse transcribe mRNA into cDNA. Following this, quantitative realtime PCR (qRT-PCR) was performed using a Power SYBR Green PCR Master Mix Kit (Applied BioSystems, Inc). The primer sequences for the genes used in this study are listed in Supplemental Table 1.

*Micro-computed tomography (microCT) analysis*

MicroCT analysis (microCT 35, Scanco Medical AG, Brüttisellen, Switzerland) was performed at 6 µm isotropic voxel size as described previously ^16^. At the femoral midshaft, a total of 100 slices located 4.8-5.4 mm away from the distal growth plate were acquired for trabecular bone and the cortical bone analyses by visually drawing the volume of interest (VOI) separately. In the L4/L5 vertebrae, the region (total about 300 slices) 50 slices away from the top and bottom end plates was acquired for trabecular bone analysis. The trabecular bone tissue within the VOI was segmented from soft tissue using a threshold of 487.0 mgHA/cm^3^ and a Gaussian noise filter (sigma=1.2, support=2.0). The cortical bone tissue was using a threshold of 661.6 mgHA/cm^3^ and a Gaussian noise filter (sigma=1.2, support=2.0). Three-dimensional standard microstructural analysis was performed to determine the geometric trabecular bone volume/total volume (BV/TV) fraction, connectivity density (Conn-Dens), trabecular thickness (Tb.Th), trabecular separation (Tb.Sp), trabecular number (Tb.N), and structure model index (SMI). For analysis of cortical bone, periosteal perimeter (Ps.Pm), endosteal perimeter (Ec.Pm), porosity, cortical bone area (Ct.Ar), cortical thickness (Ct.Th), polar moment of inertia (pMOI), and tissue mineral density (TMD) were recorded.

*Hematopoietic phenotyping of bone marrow cells*

Peripheral blood of mice was collected retro-orbitally. To analyze the peripheral blood of mice, red blood cells were first lysed and then stained for myeloid (rat anti-Gr-1 APC-Cy7, BD, 557661, rat anti-Mac-1 APC, eBioscience, 17-0112-83) or lymphoid lineages (rat anti-B220 FITC, eBioscience, 11-0452-82, hamster anti-CD3 PE-Cy7, eBioscience, 25-0031-82).

Bone marrow was flushed from femurs and cellularity was quantified with 3% acetic acid in methylene blue (STEMCELL). The lineage cell compartment of the bone marrow was analyzed by staining for myeloid and lymphoid lineages as in the peripheral blood. The HSPC compartment was analyzed by staining for Lineage (biotin-Ter-119, -Mac-1, -Gr-1, -CD4, -CD8α, -CD5, -CD19 and -B220 (eBioscience, 13-5921-85, 13-0051-85, 13-5931-86, 13-0112-86, 13-0452-86, 13-0041-86, 13-0081-86, 13-0193-86) followed by staining with streptavidin-PE-TexasRed (Invitrogen, SA1017), rat anti-cKit APC-Cy7 (eBioscience, 47-1171-82), rat anti-Sca1 PerCP-Cy5.5 (eBioscience, 45-5981-82), hamster anti-CD48 APC (eBioscience, 17-0481-82) and rat anti-CD150 PE-Cy7 (Biolegend, 115914). All flow cytometry analysis was performed on a BD LSR Fortessa flow cytometer and was analyzed on FlowJo v10.5.3 for MAC.

*Statistics*

Data are expressed as means ± standard error (SEM) and analyzed by t-tests or two-way ANOVA with a bonferroni’s post-test for multiple comparisons using Prism software (GraphPad Software, San Diego, CA). For cell culture experiments, observations were repeated independently at least three times with a similar conclusion, and only data from a representative experiment are presented. Values of p<0.05 were considered significant.

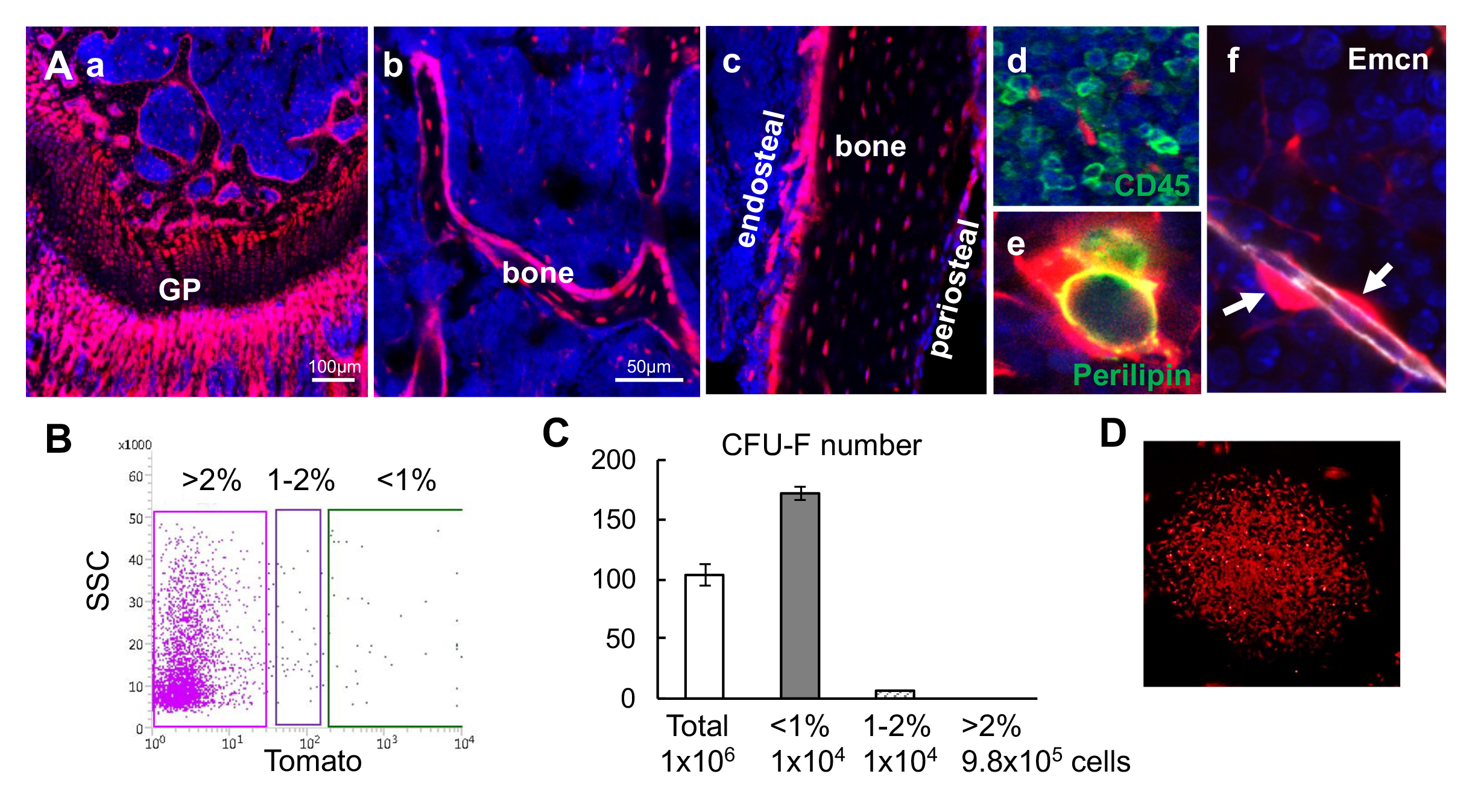

Figure S1. Bone marrow Td^+^ cells from *Col2/Td* mice contain the entire set of mesenchymal lineage cells.

(A) Fluorescent images of distal femur show that chondrocytes (a), osteoblasts and osteocytes in trabecular bone (b) and cortical bone (c), CD45^-^ stromal cells (d), adipocytes (e), and pericytes (arrows, f) are Td^+^. GP: growth plate; Emcn: Endomucin.

(B) Endosteal bone marrow cells were FACS sorted into top 1% (<1%), 1-2%, and >2% Td expressing cells.

(C) Unsorted and sorted cells were cultured for CFU-F assay. Cell number per flask is listed below. n=3-4 flasks/group.

(D) All CFU-F colonies from top 1% group were Td^+^.

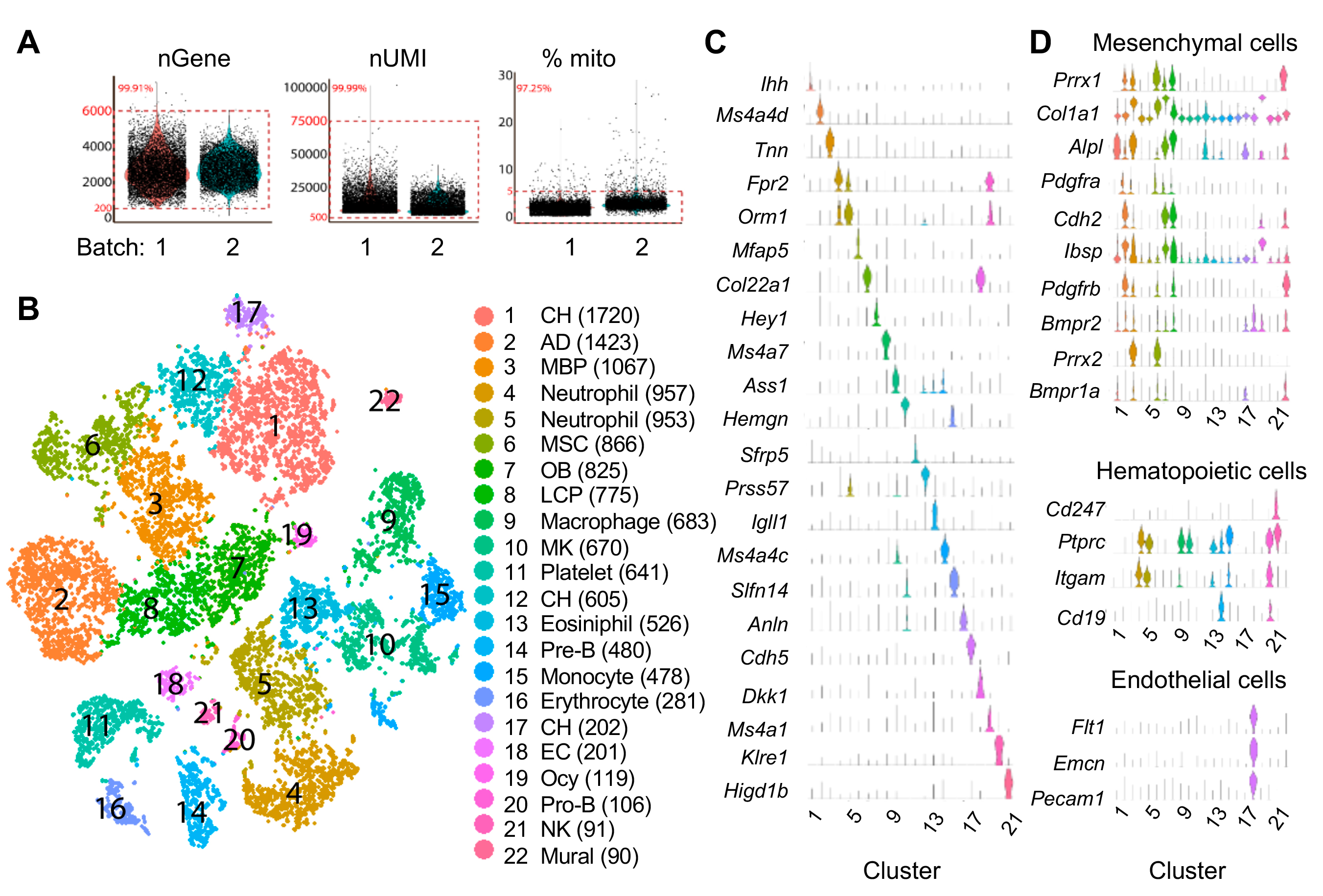

Figure S2. Large scale scRNA-seq analysis of top 1% Td^+^ cells from endosteal bone marrow of 1-1.5-month-old *Col2/Td* mice.

(A) Violin plots show numbers of genes, UMIs, and percentages of mitochondrial genes among two batches of samples (1 and 1.5 month). Red box indicates cells within the selection criteria of quality controls.

(B) The tSNE plot of 13759 Td^+^ endosteal bone marrow cells isolated from 1-1.5-month-old *Col2/Td* mice (n=5). Cell numbers are listed in parenthesis next to cluster names. MBP: mesenchymal bi-potent progenitor; OB: osteoblast; Ocy: osteocyte; LCP: lineage committed progenitor; AD: adipocyte; CH: chondrocyte; MK: megakaryocytes; EC: endothelial cells.

(C) Violin plots of cluster-specific makers.

(D) Violin plots of mesenchymal, hematopoietic, and endothelial lineage specific markers.

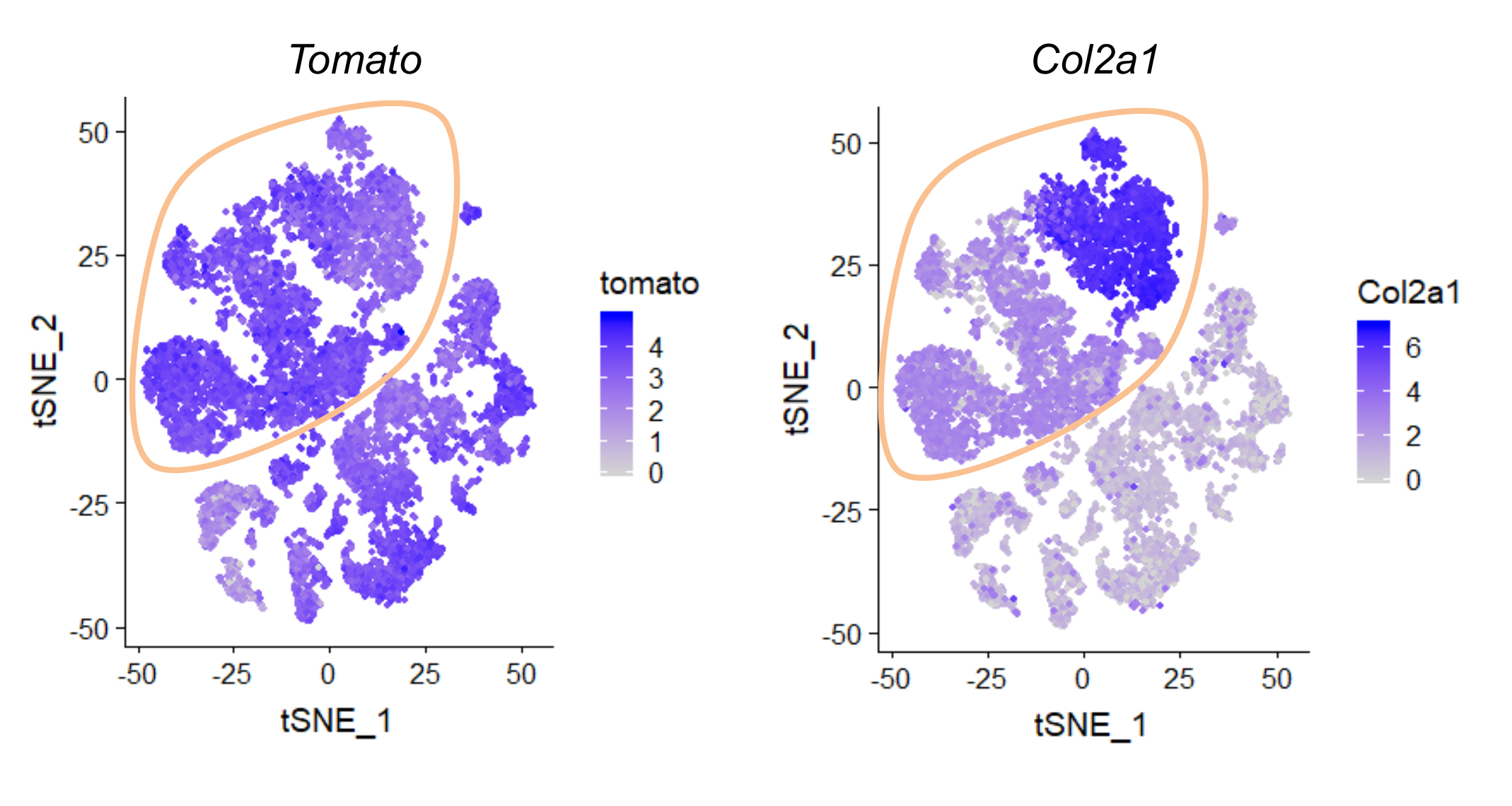

Figure S3. The expression patterns of *Tomato* and *Col2a1* in tSNE plots.

Mesenchymal lineage cells were circled. Color bar on the right of each panel indicates the expression level.

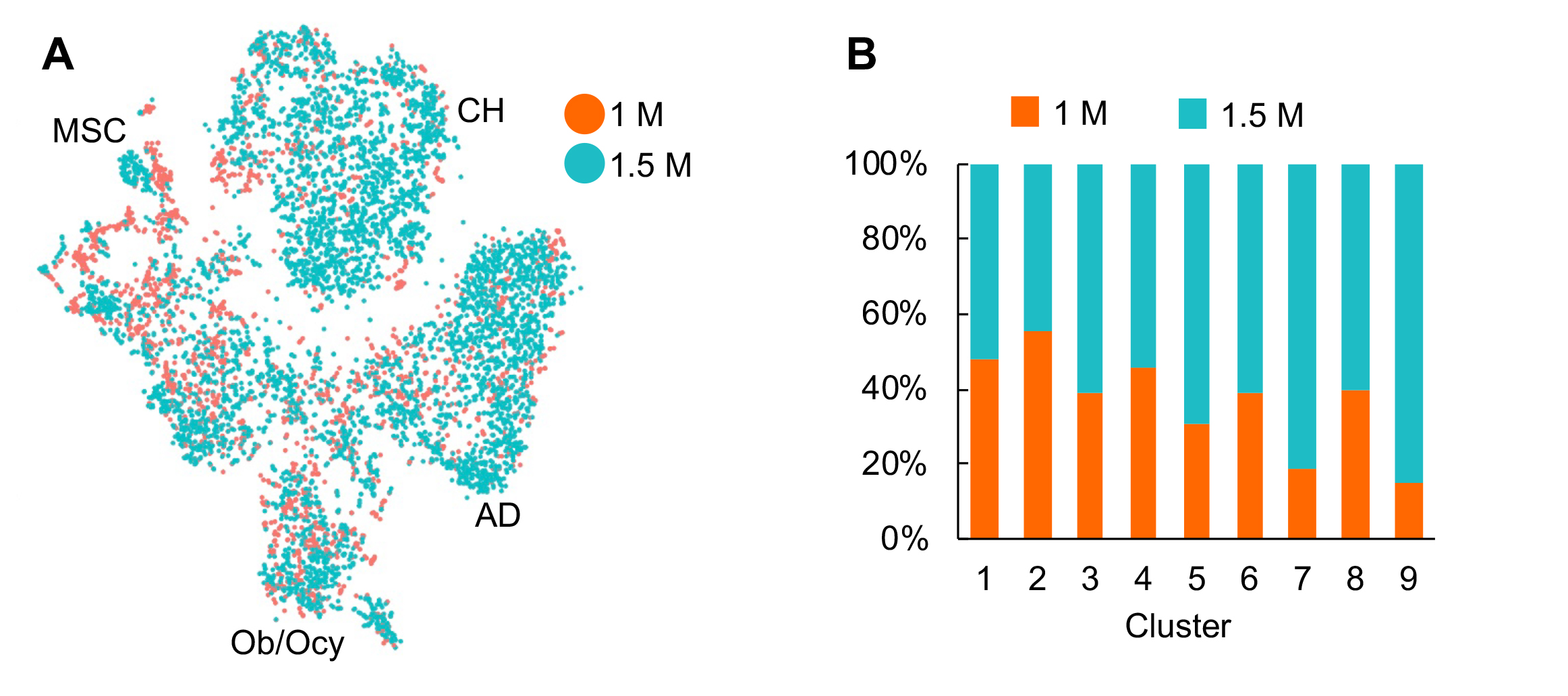

Figure S4. No batch effect was detected in our analysis.

(A) The distribution of cells from 1 month and 1.5 month datasets in the tSNE plot of mesenchymal lineage cells.

(B) The distribution of cells from 1 month and 1.5 month datasets in each cluster.

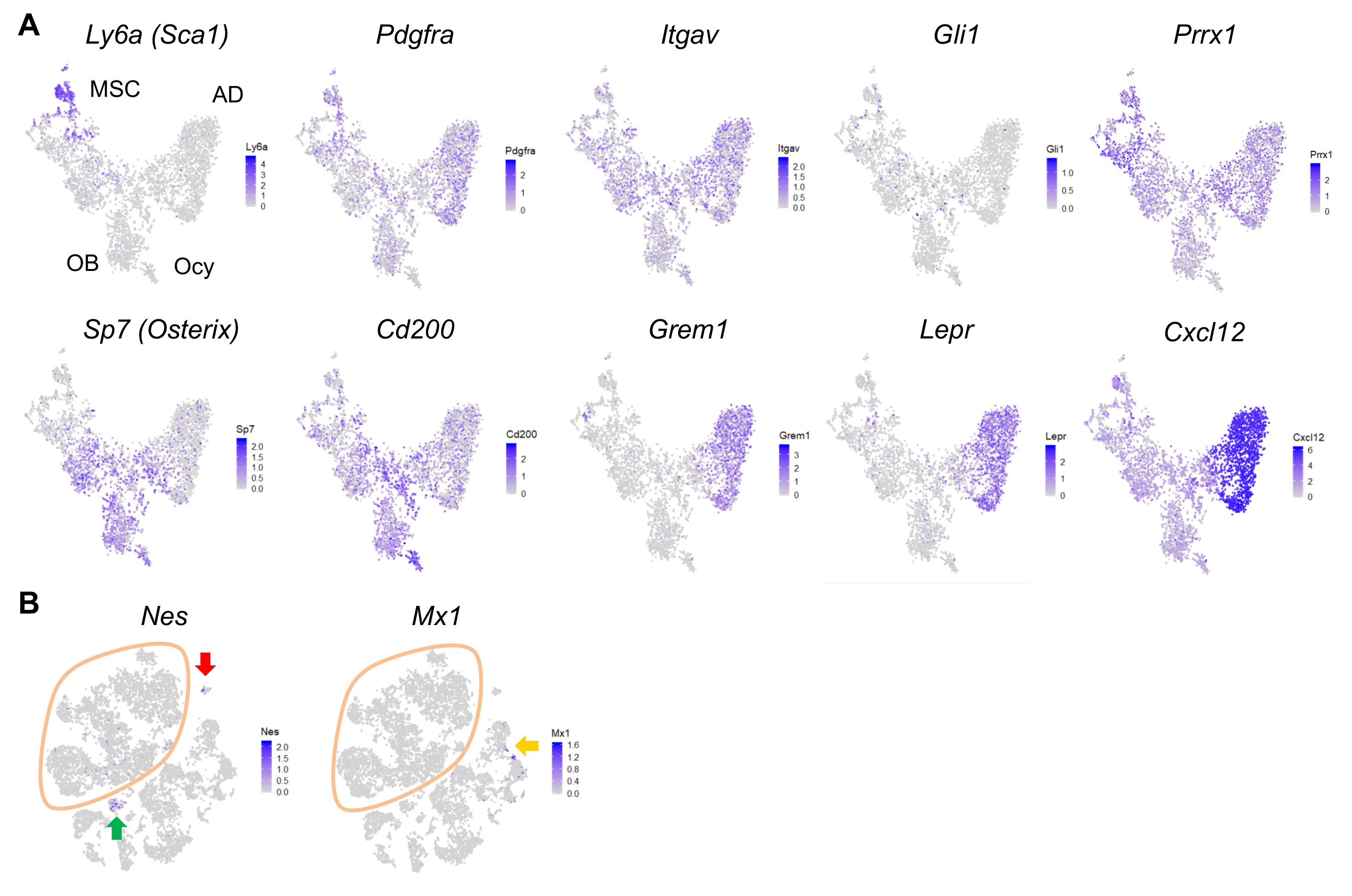

Figure S5. The expression patterns of previously reported MSC markers.

(A) The expression patterns are shown in tSNE plot that only contain mesenchymal lineage cells.

(B) The expression patterns are shown in tSNE plot that contain all sequenced cells. Green, red, and yellow arrows point to endothelial, mural, and hematopoietic cell clusters, respectively.

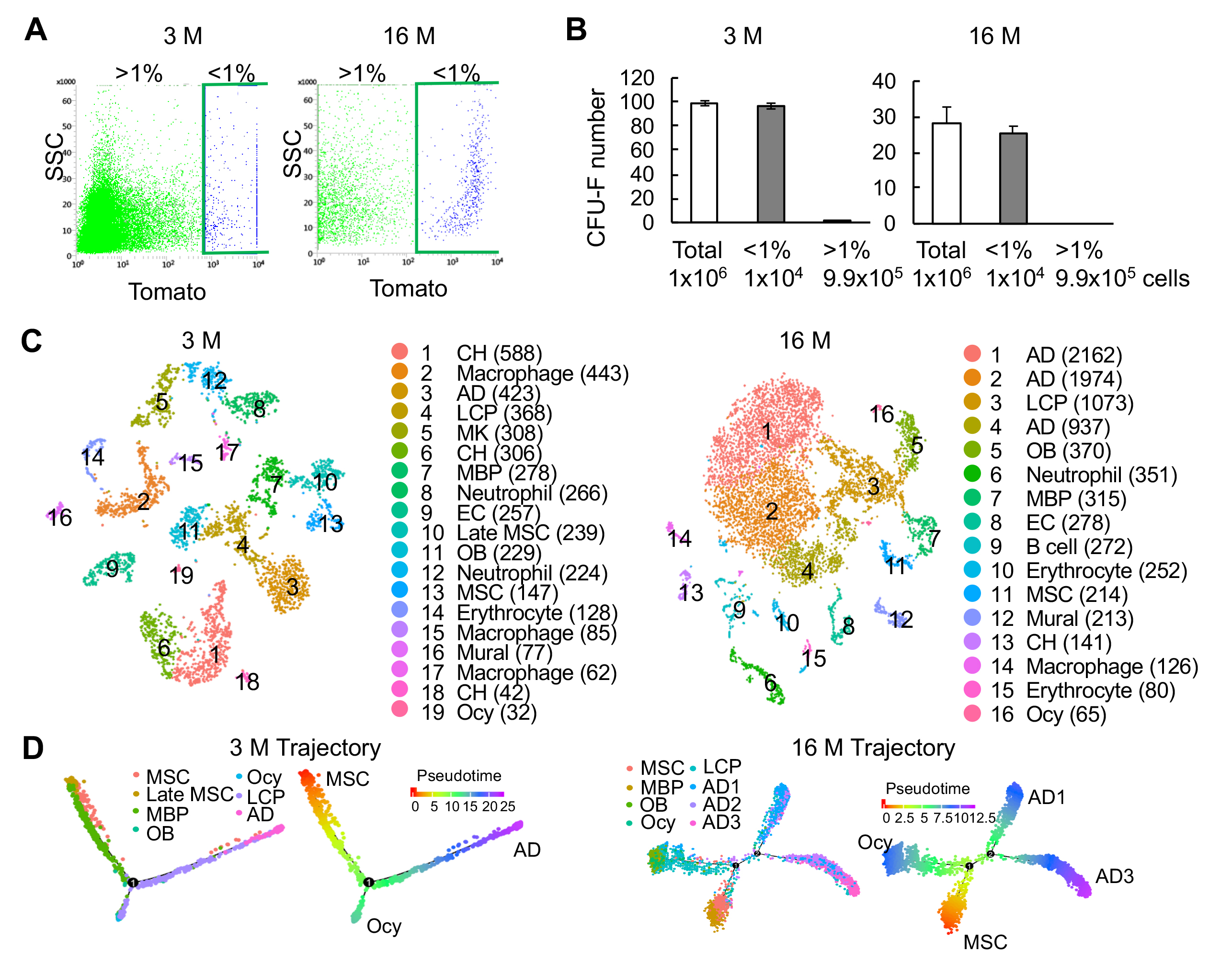

Figure S6. Large scale scRNA-seq analysis of top 1% Td^+^ cells from the endosteal bone marrow of 3 and 16-month-old *Col2/Td* mice.

(A) Endosteal bone marrow cells from 3- and 16-month old mice were FACS sorted into top 1% (<1%) and >1% Td expressing cells.

(B) Unsorted and sorted cells were cultured for CFU-F assay. Cell number per flask is listed below. n=3-4 flasks/group.

(C) The tSNE plot of 4502 and 8823 Td^+^ endosteal bone marrow cells isolated from 3- and 16-month-old *Col2/Td* mice, respectively (n=3/age). Cell numbers are listed in parenthesis next to cluster names. MBP: mesenchymal bi-potent progenitor; OB: osteoblast; Ocy: osteocyte; LCP: lineage committed progenitor; AD: adipocyte; CH: chondrocyte; MK: megakaryocyte; EC: endothelial cell.

(D) Monocle trajectory plot of bone marrow mesenchymal lineage cells (excluding chondrocytes) in 3 and 16 month datasets. Cells are labeled according to their Seurat clusters. Pseudotime scale is shown on the right.

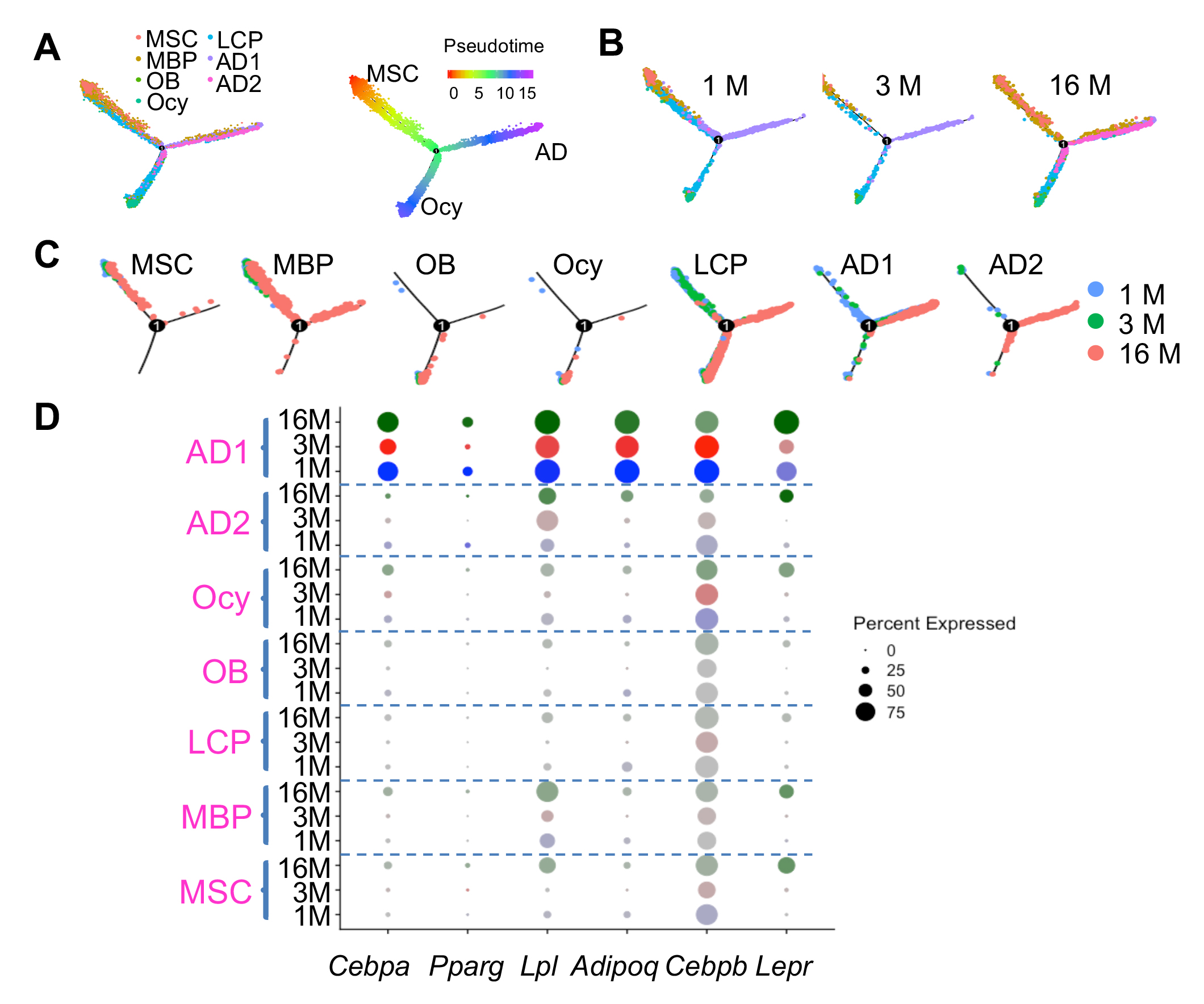

Figure S7. Pseudotime trajectory analysis of combined bone marrow mesenchymal lineage cells from all age groups.

(A) Monocle trajectory plot of bone marrow mesenchymal lineage cells of integrated database from mice at different age (1, 3 and 16 months). Cells are labeled according to their Seurat clusters. Pseudotime scale is shown on the right.

(B) Monocle trajectory plots are separated based on age groups.

(C) Monocle trajectory plots are separated based on age groups and Seurat clusters.

(D) Dotplot of *Cebpa*, *Pparg*, *Lpl*, *Adipoq* and *Lepr* in Seurat clusters across different age groups. The circle size is proportional to the percentage of cells expressing the gene and transparency of circle is reversely correlated with the average gene expression level.

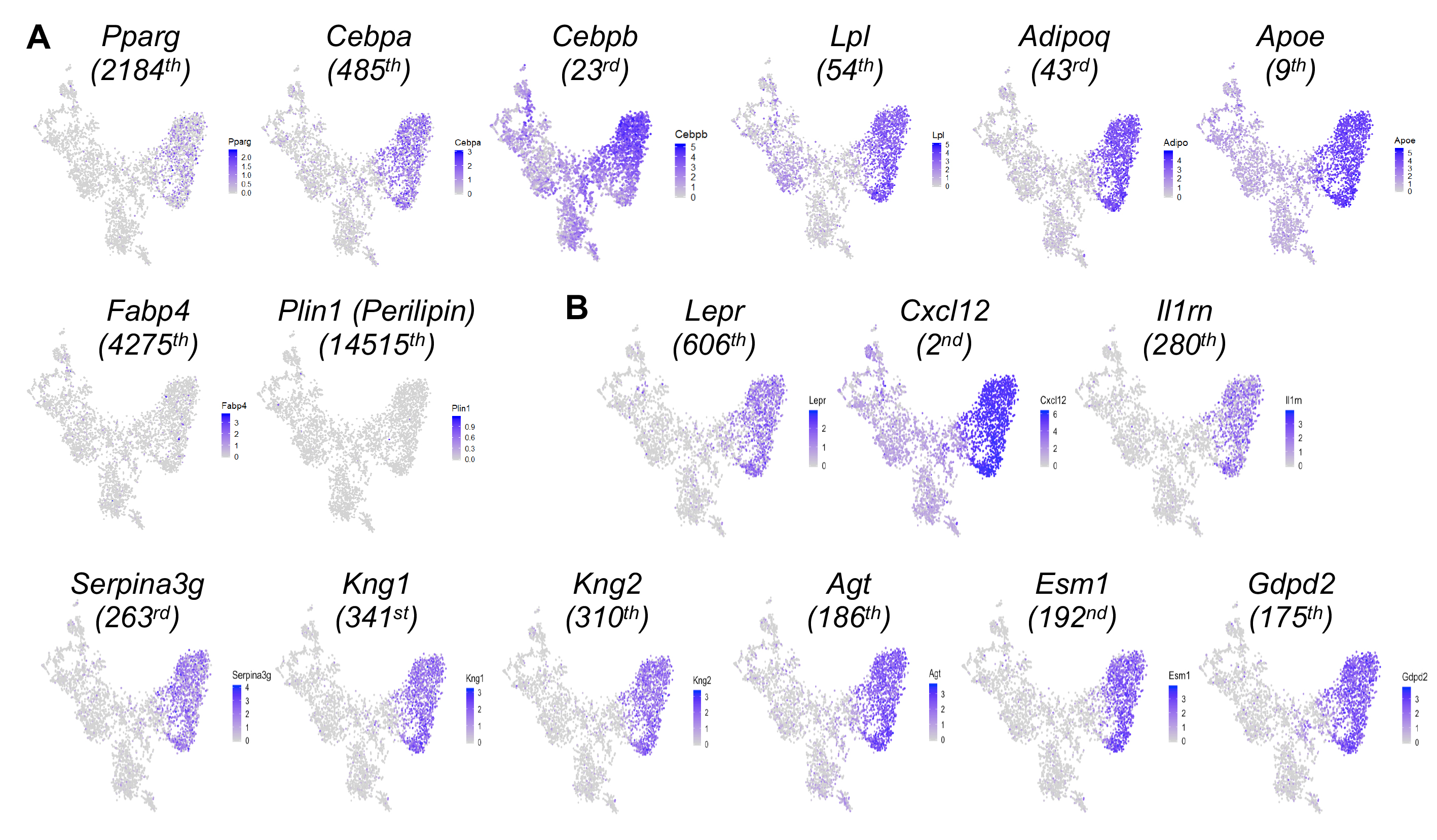

Figure S8. Large scale scRNA-seq data predict a large cluster of adipocytes in the bone marrow of adolescent mice.

(A) The expression patterns of known adipocyte markers.

(B) The expression patterns of predicted markers for the adipocyte cluster.

The number in parenthesis below each gene indicates its rank among all genes based on its expression level in adipocyte clusters.

 
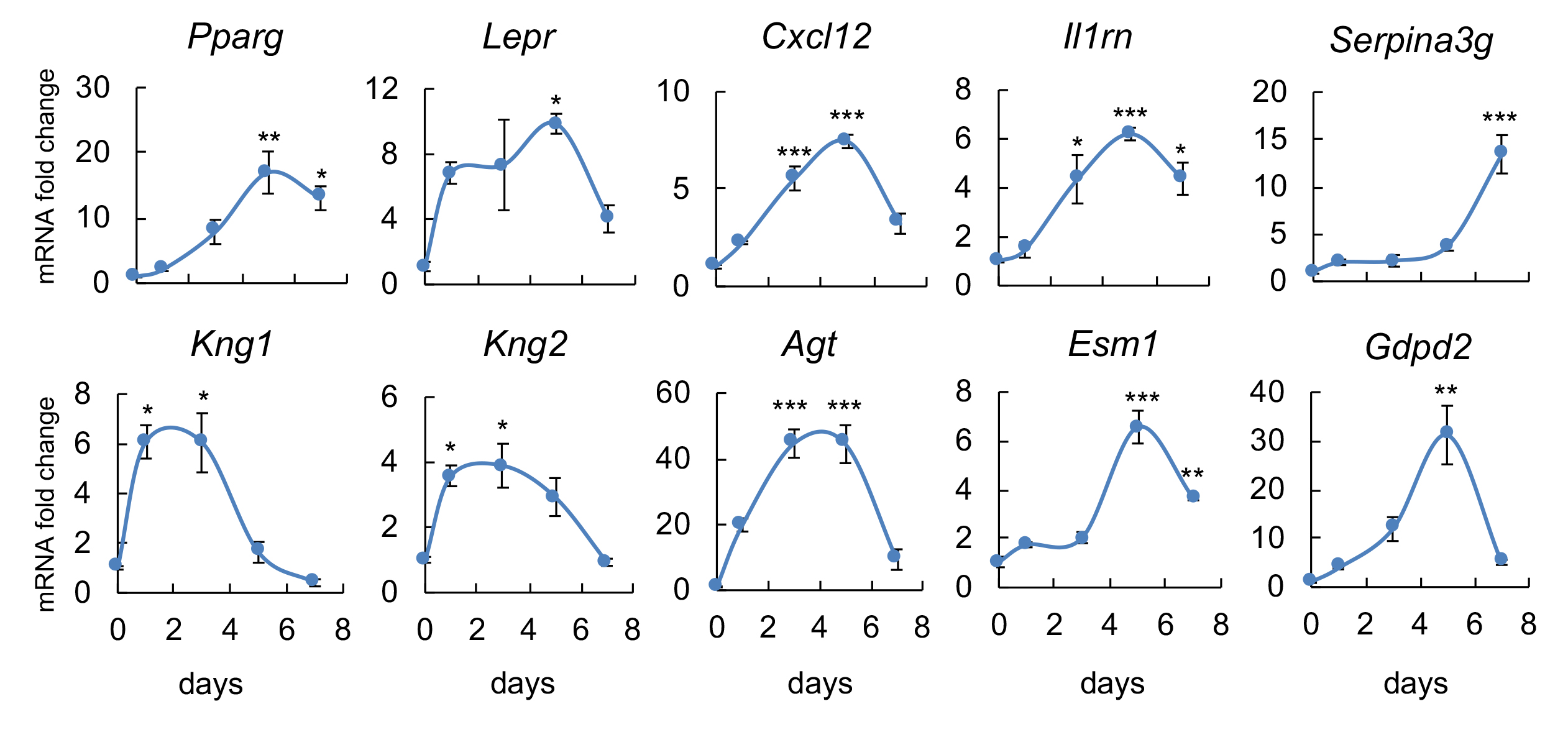

Figure S9. The adipocyte markers predicted by sequencing data are validated by in vitro adipogenic differentiation assay.

Bone marrow mesenchymal progenitors were cultured to confluence and switched to adipogenic differentiation medium. Cells were harvested at indicated time points for qRT-PCR analysis of newly identified marker genes. *: p<0.05; **: p<0.01; ***: p<0.001 compared with day 0.

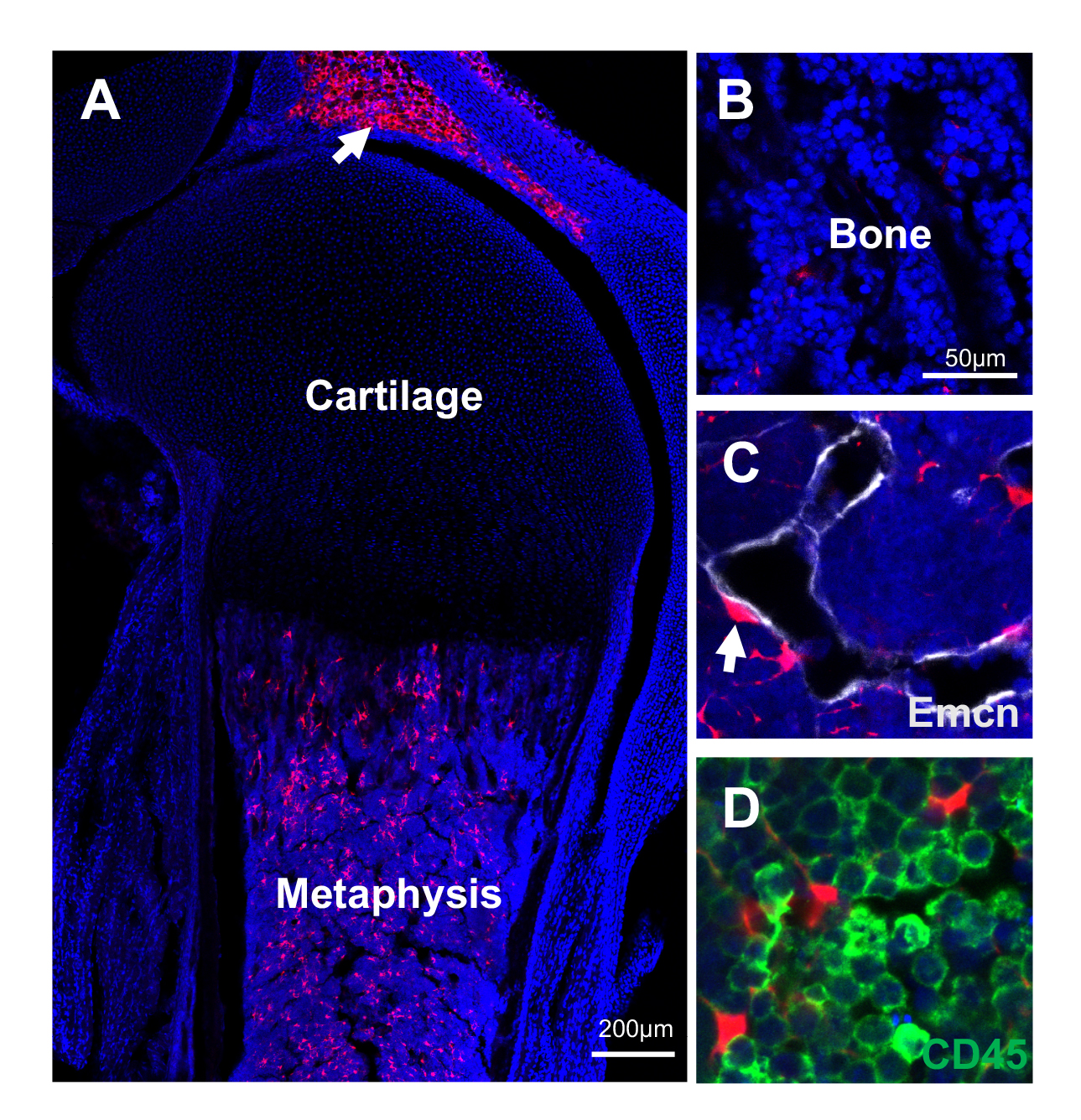

Figure S10. *Adipoq/Td* mice at P5 have abundant Td^+^ adipocytes in the bone marrow.

(A) Fluorescent image of the proximal tibia at a low magnification. Arrow points to patellar fat pad.

(B-D) At a high magnification, it is obvious that Td does not label osteoblasts and osteocytes in trabecular bone (B) but does label pericytes (C, arrow) and CD45^-^ stromal cells (D).

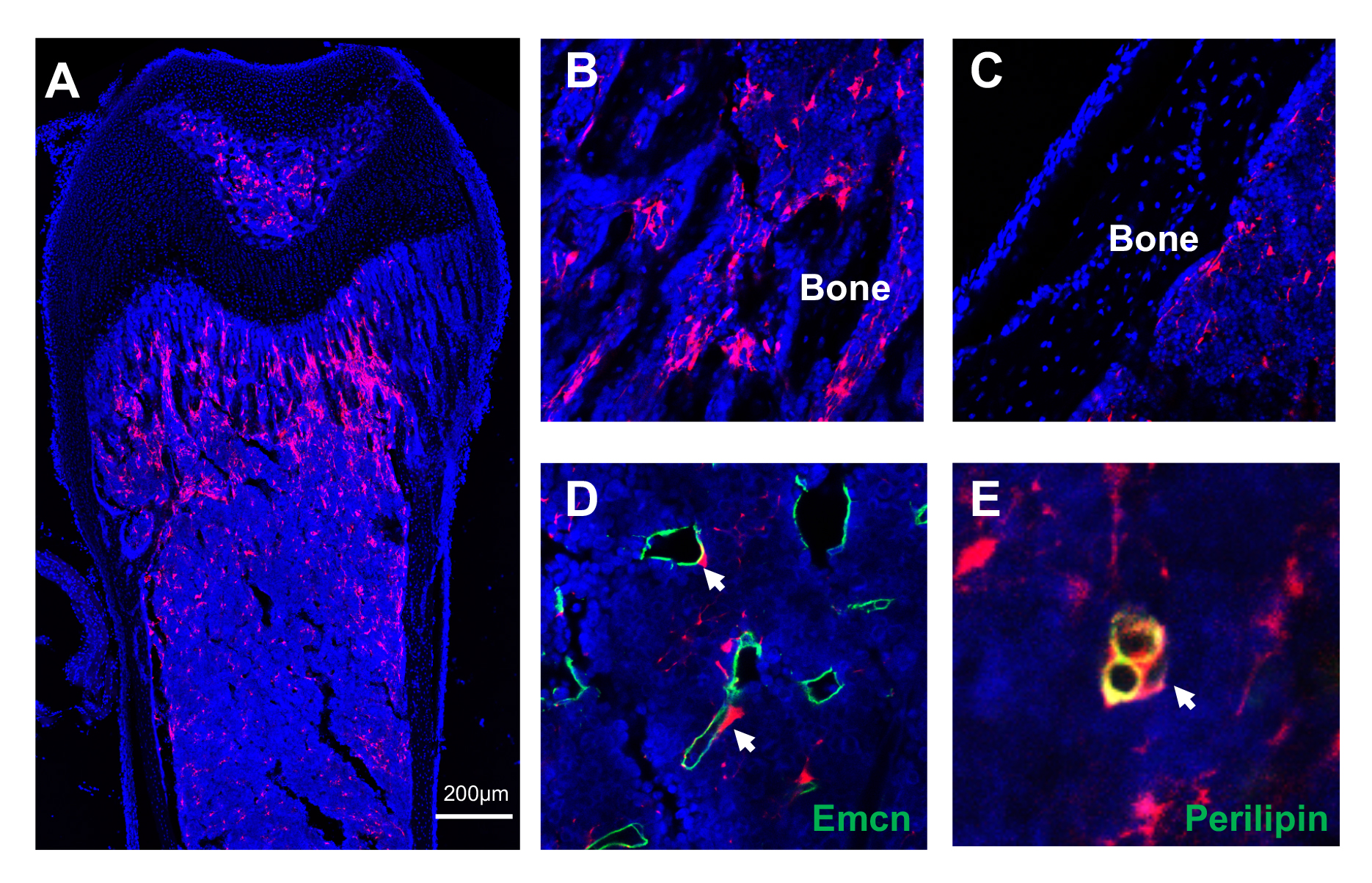

Figure S11. Young *AdipoqER/Td* mice have abundant bone marrow Td^+^ cells.

(A) Low magnification image of a femur from a 1-month old *AdipoqER/Td* mouse (Tamoxifen at 2 weeks) reveals abundant Td^+^ cells in the bone marrow. Td does not label chondrocytes in articular cartilage and growth plate.

(B-E) High magnification images show that Td^+^ cells are located inside bone marrow (B). Similar to *Adipoq/Td* mice, Td^+^ cells in *AdipoqER/Td* mice are not osteoblasts and osteocytes (C). They are indeed pericytes (arrows, D) and Perilipin^+^ adipocytes (arrow, E).

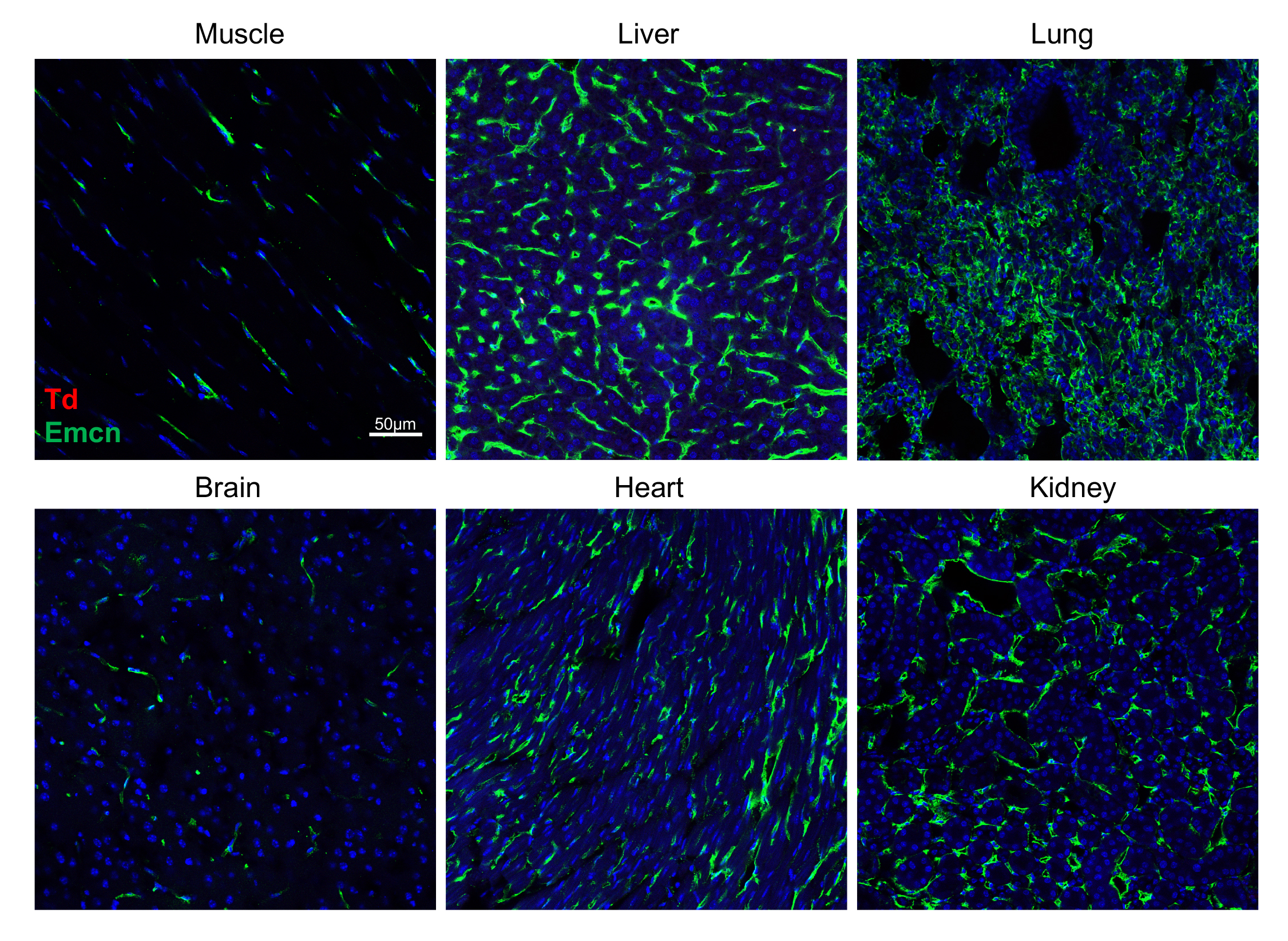

Figure S12. Td^+^ pericytes are unique to the bone marrow of *Adipoq/Td* mice.

Td^+^ pericytes are not detected in muscle, liver, lung, brain, heart, and kidney of 1-month-old *Adipoq/Td* mice. Emcn: vessels. n=3 mice.

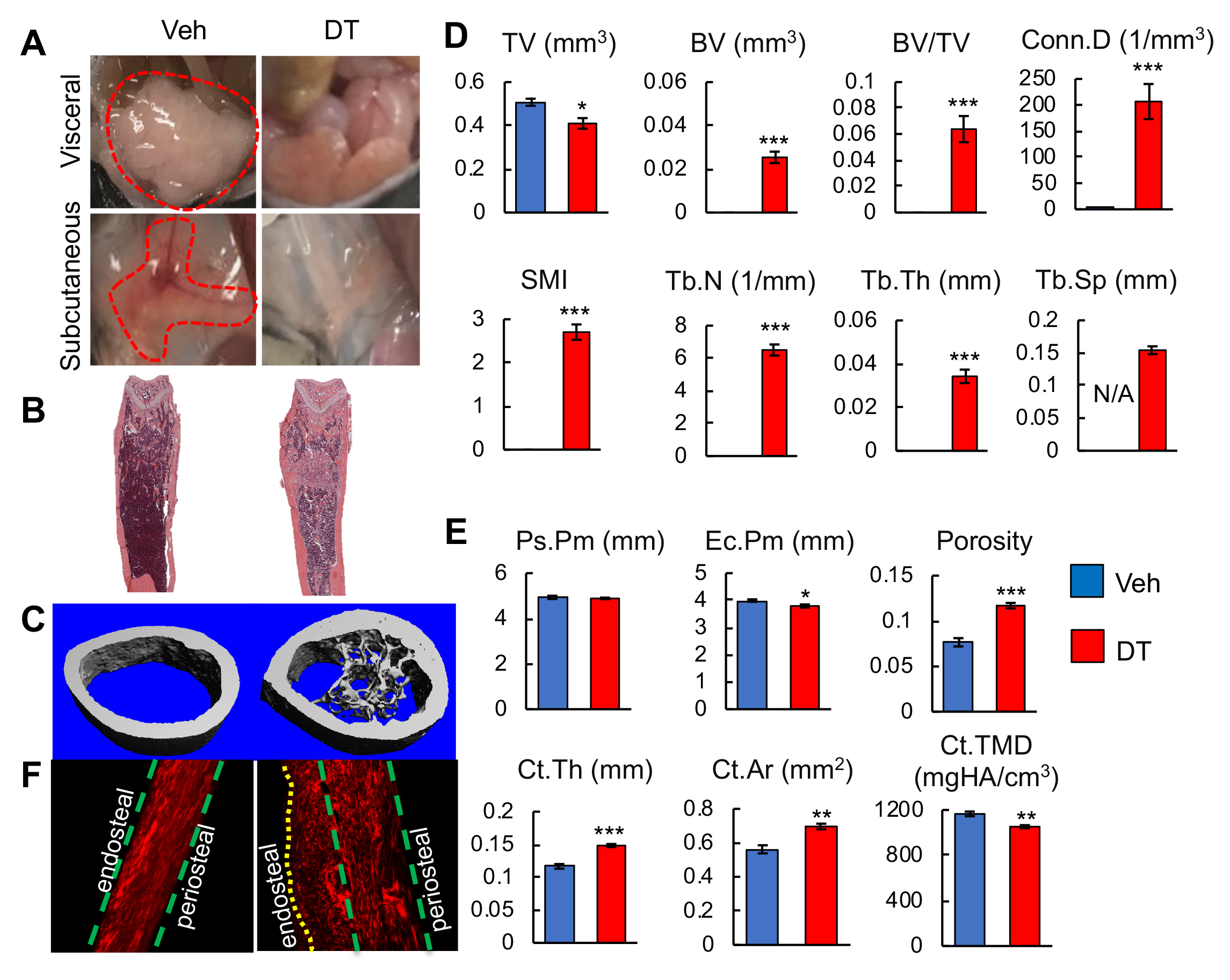

Figure S13. Ablation of bone marrow non-lipid-laden adipocytes rapidly stimulates new bone formation in the long bone diaphyseal area.

(A) 2 weeks of DT injections significantly reduce the volume of visceral and subcutaneous fat tissues (dash line) in 1-month-old *Adipoq/Td/DTR* mice.

(B) H&E staining of femoral sections from *Adipoq/Td/DTR* mice at 2 weeks after vehicle or DT injections.

(C) 3D reconstructed microCT images of femoral midshaft region.

(D) MicroCT measurement of trabecular bone structural parameters from the midshaft region.

(E) MicroCT measurement of cortical bone structural parameters from the midshaft region.

n=6 mice/group. *: p<0.05; **: p<0.01; ***: p<0.001 DT vs veh.

(F) Second harmonic generation images of femoral sections reveal that collagen fibers in the newly formed bone on the endocortical surface are mostly misaligned in *Adipoq/Td/DTR* mice after DT injections.

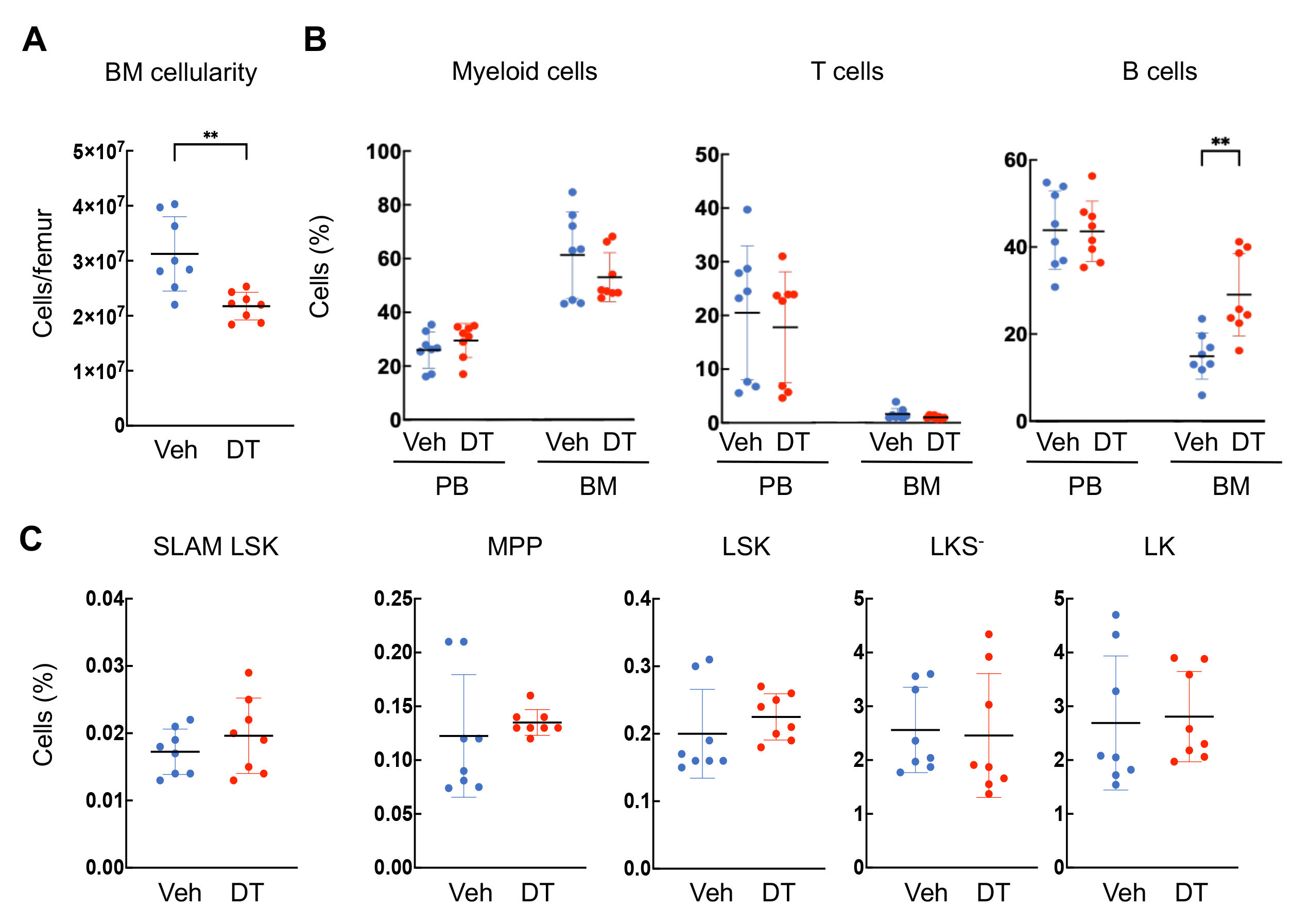

Figure S14. Ablation of bone marrow non-lipid-laden adipocytes reduces overall bone marrow (BM) cellularity but has little effect on hematopoietic cells.

(A) 2 weeks of DT injections significantly reduce femoral BM cellularity in 1-month-old *Adipoq/Td/DTR* mice.

(B) Peripheral blood (PB) and BM frequency of hematopoietic lineage cells were assessed by flow cytometry.

(C) Bone marrow frequency of various hematopoietic stem and progenitor cell compartments were assessed by flow cytometry using SLAM marker scheme. Hematopoietic stem cell (SLAM LSK): Lin^-^Sca1^+^cKit^+^CD48^-^CD150^+^; Multipotent progenitor (MPP): Lin^-^Sca1^+^cKit^+^CD48^+^CD150^-^. LSK: Lin^-^Sca1^+^cKit^+^; LKS^-^: Lin^-^Sca1^-^cKit^+^; LK: Lin^-^cKit^+^.

Each symbol represents an individual mouse.

Horizontal lines represent the mean and vertical bars indicate the SD. **: p<0.01 DT vs veh.

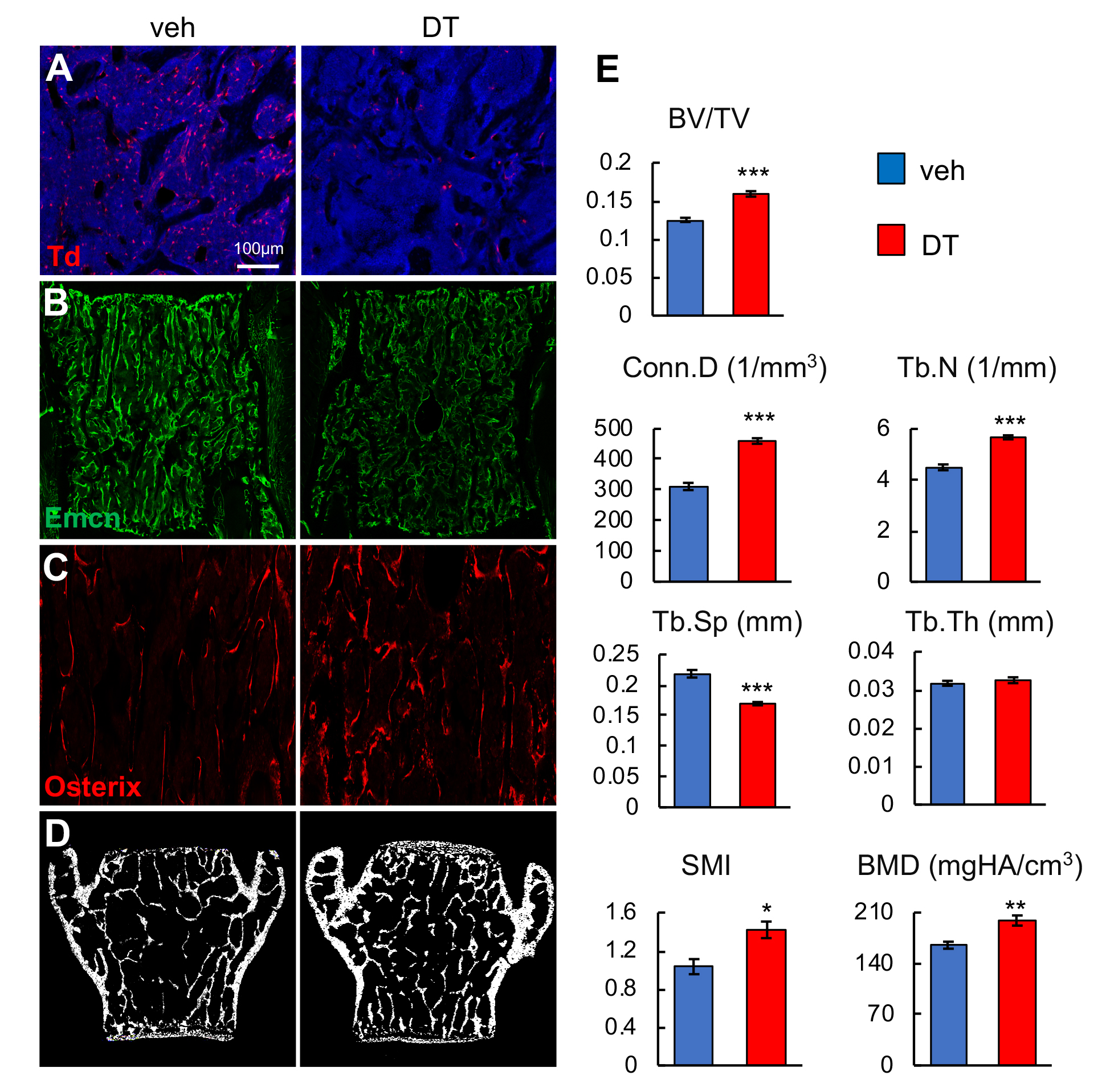

Figure S15. Ablation of bone marrow non-lipid-laden adipocytes also increases trabecular bone mass in vertebrae.

(A) 2 weeks of DT injections significantly reduce the number of bone marrow Td^+^ cells in vertebrae of 1-month-old *Adipoq/Td/DTR* mice.

(B) Vessel staining reveals that DT injections alter vessel structure and integrity in the vertebral bone marrow of *Adipoq/Td/DTR* mice.

(C) Osterix staining suggests more bone forming cells after adipocyte ablation.

(D) Representative microCT images of vertebral bone.

(E) MicroCT measurement of trabecular bone structural parameters in vertebrae. n=6 mice/group. *: p<0.05; **: p<0.01; ***: p<0.001 DT vs veh.

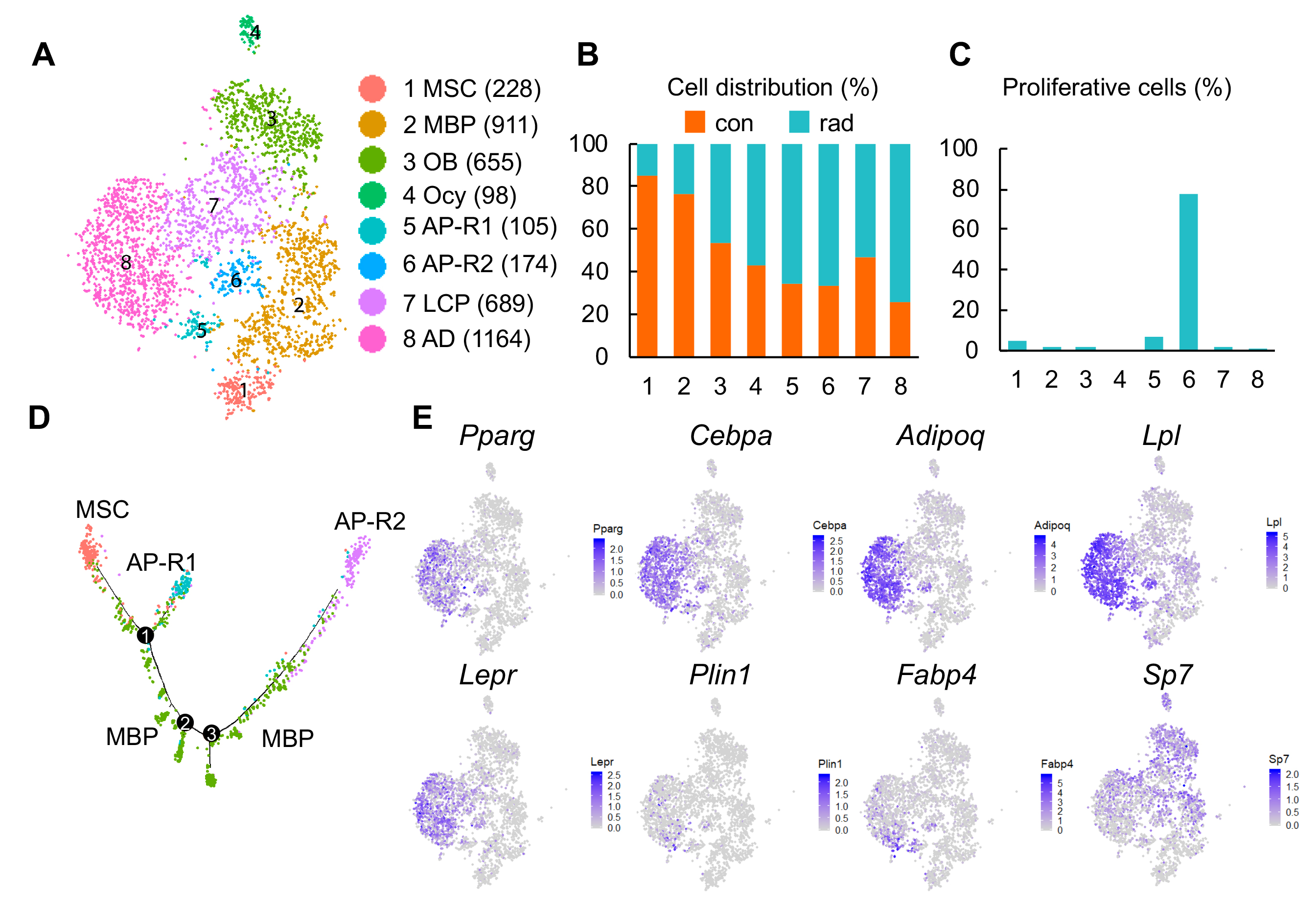

Figure S16. Radiation generates new adipogenic differentiation routes from mesenchymal progenitors.

(A) TSNE plot of endosteal mesenchymal lineage cells from control and radiated animals. Cell numbers are listed in parenthesis next to cluster names. MBP: mesenchymal bi-potent progenitor; OB: osteoblast; Ocy: osteocyte; LCP: lineage committed progenitor; AD: adipocyte; AP: adipoprogenitor.

(B) The distribution of control and radiated mesenchymal lineage cells in each cluster.

(C) The percentage of proliferative cells (S/G2/M phase) in each cluster.

(D) The pseudotime plot of cells in MSC, MBP, AP-R1, and AP-R2 clusters.

(E) The expression patterns of adipocyte markers and an osteoblast marker (Sp7) are shown in tSNE plot.

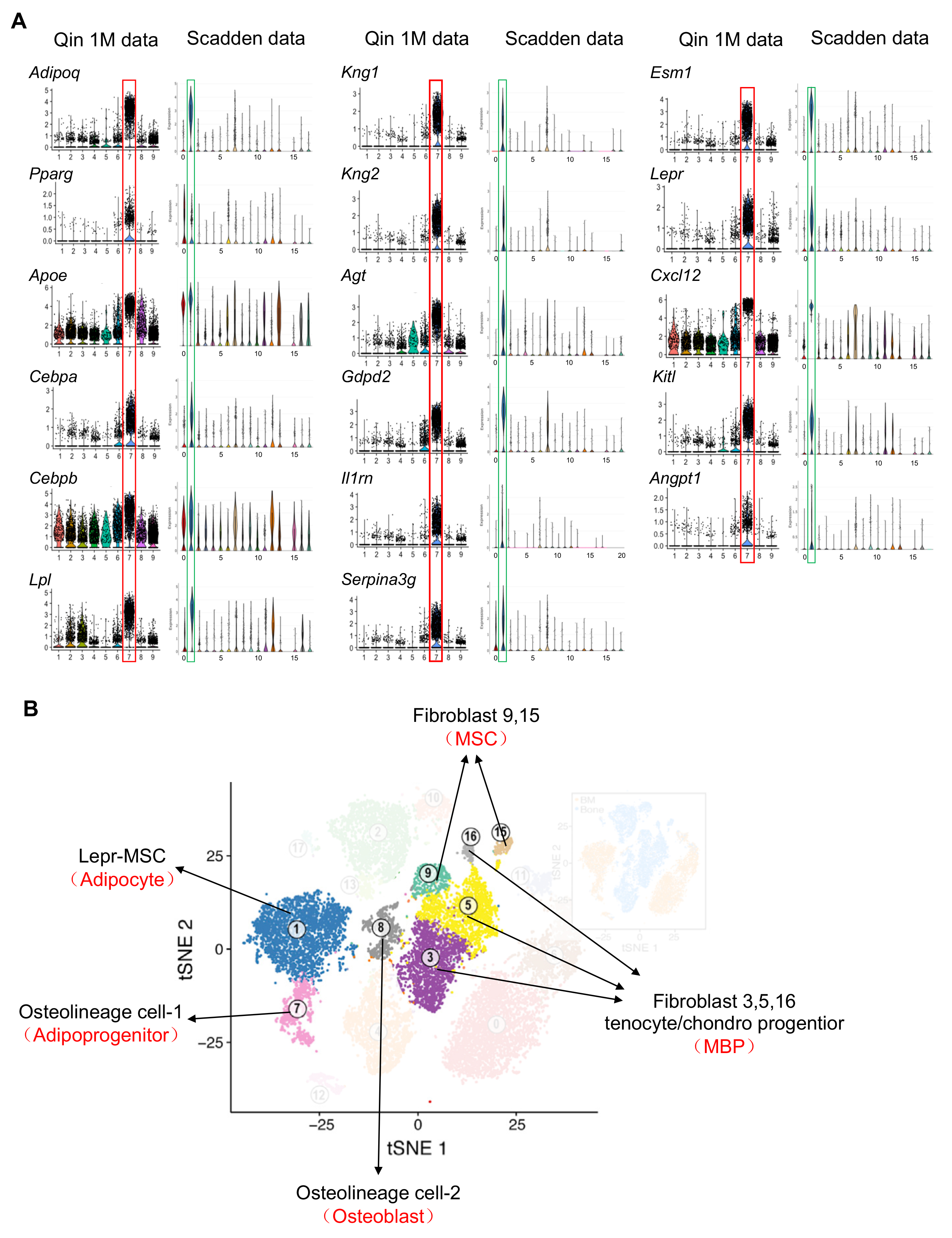

Figure S17. The comparison of large scale scRNA-seq data of bone marrow mesenchymal lineage cells between our group (Qin) and Scadden group that was recently published ^17^ confirms the same distribution of mesenchymal subpopulations but with different annotation of cell clusters.

(A) Violin plots of MERA cluster markers, including known adipocyte markers and novel markers in Qin’s 1-1.5-month-old dataset and Scadden’s dataset (available from https://portals.broadinstitute.org/single_cell/study/mouse-bone-marrow-stroma-in-homeostasis). Red boxes are cluster 7 adipocyte in Qin dataset and blue boxes are cluster 1 MSC in Scadden dataset. All those markers have the same expression patterns in adipocytes annotated by Qin group and MSCs annotated by Scadden group.

(C) An alternative annotation of cell clusters in the tSNE plot of Scadden dataset. Our annotation results, indicated by red in parentheses below Scadden’s annotation, are based on examining violin plots of all our cluster marker genes in Scadden dataset.

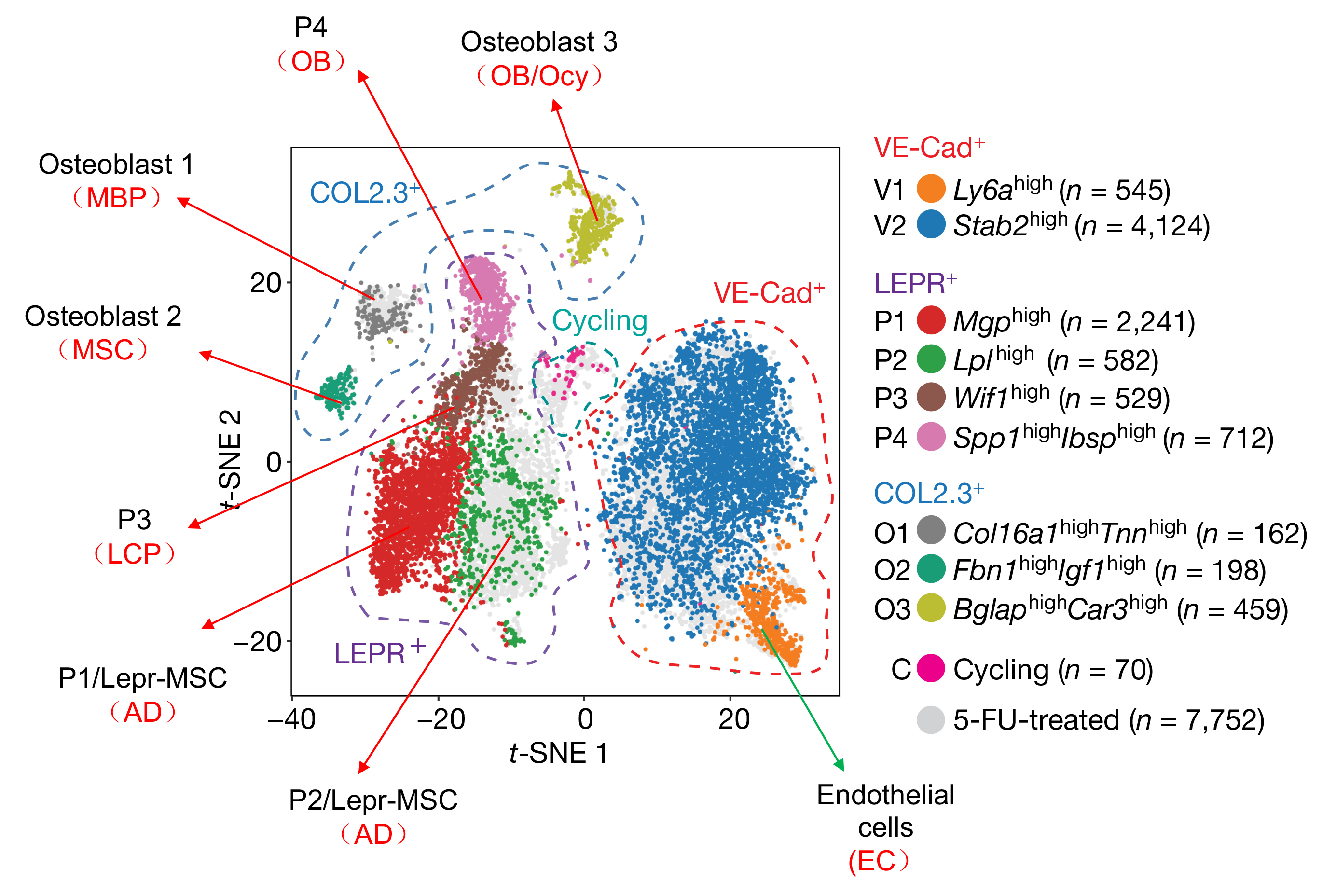

Figure S18. A comparison of large scale scRNA-seq data of bone marrow mesenchymal lineage cells between our group and Aifantis group that was recently published ^18^. An alternative annotation of cell clusters based on our data are indicated by red in parentheses below Aifantis’s annotation.

**Table S1.** Mouse real-time PCR primer sequences used in this study.

| Gene | Forward primer | Reverse primer |
| --- | --- | --- |
| *Cebpa* | 5’-CAAGAACAGCAACGAGTACCG-3’ | 5’-GTCACTGGTCAACTCCAGCAC |
| *Lpl* | 5’-GGGAGTTTGGCTCCAGAGTTT-3’ | 5’-TGTGTCTTCAGGGGTCCTTAG |
| *Adipoq* | 5’-AAAGGAGAGCCTGGAGAA-3’ | 5’-GAATGGGTACATTGGGAACA-3’ |
| *Pparg* | 5’-CCAGCGTGAAGCCAGAGTAG-3’ | 5’-ACCGTGGCTGTGCTCATCCT-3’ |
| *Lepr* | 5’-TGGTCCCAGCAGCTATGGT-3’ | 5’-ACCCAGAGAAGTTAGCACTGT-3’ |
| *Cxcl12* | 5’-CTGTGCCCTTCAGATTGTT-3’ | 5’-AGCTTTCTCCAGGTACTCTT-3’ |
| *Serpina3g* | 5’-CTGTGGTGGAGCTGAAATAC-3’ | 5’-TCAGGGTCTCTGGTTGTAAG-3’ |
| *Agt* | 5’-TCCACTGACCCAGTTCTT-3’ | 5’-AAGTAGGGTGCTGTCTGT-3’ |
| *Gdpd2* | 5’-CAGGAGTGGCATAGTTTACG-3’ | 5’-CAGGCCAACAAGGTGATT-3’ |
| *Il1rn* | 5’-TGCCAAGTCTGGAGATGA-3’ | 5’-CAGAGCGGATGAAGGTAAAG-3’ |
| *Kng1* | 5’-GAGACCTTGGGAGAACAAAG-3’ | 5’-ACACTCCGGAAAGGAGAA-3’ |
| *Kng2* | 5’-GGACTCCTGCTGACTTTAAC-3’ | 5’-GCATCCACAGCCTGAAATA-3’ |
| *Esm1* | 5’-TACAGCGAGGAGGATGATT-3’ | 5’-GGCAATTGCAAGTCTCTTTG-3’ |
| *Tnfsf11* | 5’-CAGCATCGCTCTGTTCCTGTA-3’ | 5’-CTGCGTTTTCATGGAGTCTCA-3’ |
| *Vegfa* | 5’-GCACATAGAGAGAATGAGCTTCC-3’ | 5’-CTCCGCTCTGAACAAGGCT-3’ |
| *Vegfc* | 5’-GAGGTCAAGGCTTTTGAAGGC-3’ | 5’-CTGTCCTGGTATTGAGGGTGG-3’ |
| *Rspo3* | 5’-ATGCACTTGCGACTGATTTCT-3’ | 5’-GCAGCCTTGACTGACATTAGGAT-3’ |
| *Angpt4* | 5’-CAGCCAGCTATGCTACTAGATGG-3’ | 5’-CCTCTGGAGGCTATTGGAGC-3’ |
| *Actb* | 5’-GGCTGTATTCCCCTCCATCG-3’ | 5’-CCAGTTGGTAACAATGCCATGT-3’ |
