## Supplementary material for "Single cell transcriptomics identifies a unique adipocyte population that regulates bone marrow environment": video legend

Video 1: Confocal fluorescence image of Td^+^ cells in bone marrow of *Adipoq/Td* mice to show its 3D network structure made of cell processes protruding from cell bodies. Scan depth: 50 μm.

Video 2: Confocal fluorescence image of Td^+^ cells with PDGFRβ staining (green) in bone marrow of *Adipoq/Td* mice. Scan depth: 50 μm.

Video 3: Confocal fluorescence image of Td^+^ cells with Connexin 43 staining (green, shown as dots on cell processes) in bone marrow of *Adipoq/Td* mice. Scan depth: 50 μm.

Video 4: Confocal fluorescence image of Td^+^ cells with Emcn staining (green, vessel) in bone marrow of *Adipoq/Td* mice. Scan depth: 50 μm.

Video 5: Confocal fluorescence image of Td^+^ cells with Perilipin staining (green) in bone marrow of *Adipoq/Td* mice. Large yellow circles are lipid-laden adipocytes. Scan depth: 50 μm.
